# Heritable variation in locomotion, reward sensitivity and impulsive behaviors in a genetically diverse inbred mouse panel

**DOI:** 10.1101/2021.04.06.438678

**Authors:** Lauren S. Bailey, Jared R. Bagley, Rainy Dodd, Ashley Olson, Mikayla Bolduc, Vivek M. Philip, Laura G. Reinholdt, Stacey J. Sukoff Rizzo, Leona Gagnon, Elissa J. Chesler, J. David Jentsch

**Affiliations:** State University of New York – Binghamton University; The Jackson Laboratory, Bar Harbor ME; University of Pittsburgh School of Medicine, Pittsburgh, PA

## Abstract

Drugs of abuse, including alcohol and stimulants like cocaine, produce effects that are subject to individual variability, and genetic variation accounts for at least a portion of those differences. Notably, research in both animal models and human subjects point towards reward sensitivity and impulsivity as being trait characteristics that predict relatively greater positive subjective responses to stimulant drugs. Here we describe use of the eight Collaborative Cross (CC) founder strains and multiple CC strains to examine the heritability of reward sensitivity and impulsivity traits, as well as genetic correlations between these measures and existing addiction-related phenotypes. *Methods*. Strains were all tested for activity in an open field and reward sensitivity (intake of chocolate BOOST®). Mice were then divided into two counterbalanced groups and underwent reversal learning (impulsive action and waiting impulsivity) or delay discounting (impulsive choice). *Results*. CC and founder mice demonstrate significant heritability for impulsive action, impulsive choice, waiting impulsivity, locomotor activity, and reward sensitivity, with each impulsive phenotype determined to be non-correlating, independent traits. This research was conducted within the broader, inter-laboratory effort of the Center for Systems Neurogenetics of Addiction (CSNA) to characterize CC and DO mice for multiple, cocaine abuse related traits. These data will facilitate the discovery of genetic correlations between predictive traits, which will then guide discovery of genes and genetic variants that contribute to addictive behaviors.

## Introduction

Not all individuals who initiate drug or alcohol use will eventually lose control over their use and/or meet diagnostic criteria for Substance Use Disorder (SUD). This progression is moderated by a myriad of genetic and environmental factors that interact with one another in a developmentally- and sex-dependent manner (Bevilacqua & Goldman, 2013; Bezdijan, Baker, & Tuvblad, 201; Kreek et al., 2005; Piazza & Deroche-Gamonet, 2013). Evidence from twin studies supports the idea that the majority of risk for developing an SUD relates to a single substance-nonspecific genetic factor, with lesser influences of common and unique environmental influences (Kendler et al., 2003), but more recent genome-wide association studies indicate a multitude of common and distinct genetic factors (Crist, Reiner, & Berrettini, 2019; Hancock et al., 2018). A major focus of current addiction studies is therefore to discover potential genetic and neurobiological factors that moderate individual SUD risk. Multiple heritable phenotypes have been shown to be predictive for heightened likelihood to seek out and use drugs in both humans and laboratory animals, including novelty preference and seeking (Belin et al., 2011; Molander et al., 2011; Wingo et al., 2015), locomotor response to novelty (Nadal, Armario, & Janak, 2002; Piazza & Deroche-Gamonet, 2013), anxiety-related behaviors (Giles, Turk, & Fresco, 2006; Spanagel et al., 1995; Stathopoulou et al., 2021), altered stress responses (Enoch, 2010; Kreek & Koob, 1998; Sinha, 2008), circadian phenotypes (Logan, Williams, & McClung, 2014; Rosenwasser, 2010), and impulsivity. These behaviors may have overlapping neurogenetic components, and by studying them in tandem, common and unique genetic factors may be identified and investigated for their relationship to SUD traits.

Impulsivity is the trait-like proclivity to engage in excessive, uncontrolled, or rash reward pursuit and consumption (Dalley, Everitt, & Robbins, 2011; Evenden, 1999; Jentsch et al., 2014). These behaviors are considered pathological when they are intrusive, disrupt normal life routines, cause clinical distress, or lead to harmful outcomes (Moeller et al., 2001). Impulsivity is furthermore identified as having multiple dimensions, each of which can be separately measured and may have a unique relationship to addiction vulnerability. This is exemplified in the widely used Barratt Impulsiveness Scale (BIS-11), which measures self-reported cognitive, motor, and non-planning impulsiveness (Barratt, 1959). Behavioral tasks have also been developed to quantify impulsive phenotypes in humans, such as the Go/No-Go task, delay discounting, reversal learning, Five-Choice Serial Reaction Time Task (5-CSRTT), and stop signal reaction time task, which have analogs for use in animal models. Research on animal models exposed a unique relationship between types of impulsivity and various pharmacological interventions, in that drug treatment did not uniformly increase, decrease, or maintain different impulsive phenotypes (Evenden, 1998; Evenden, 1999; Winstanley et al., 2004). No significant bivariate correlation between impulsive action (five-choice serial reaction time task) and impulsive choice (delay discounting) has been found in either lab rats or humans; and furthermore, three factors reflecting statistically orthogonal measures of impulsivity were identified in human subjects: self-report, impulsive action, and impulsive choice (Broos et al., 2012; MacKillop et al., 2016). These findings may very likely be attributed to different underlying neurogenetic mechanisms regulating each type of impulsivity.

Delay discounting, a common test of impulsive choice, was initially created to assess rats and pigeons (Evenden & Ryan, 1996), though it is now used in both human and animal subjects with variations to the methodology. Delay discounting is a paradigm established to assess an individual’s tendency to reduce (discount) the subjective value of a reward if it must wait to receive it. The delay discounting procedure aims to establish how the subject therefore discounts the delayed reward, either by altering the volume of reward or the length of delay. A fundamental aspect of delay discounting is that the subject makes an action to choose one of two options and then must wait for the consequence, thus differing from other similar procedures such as the differential reinforcement task (Evenden, 1999), and measuring impulsive choice (frequent selection of the small immediate lever) versus self-control (frequent selection of the large, delayed option) (Odum, 2013).

Reversal learning, a measure hypothetically linked with impulsive action, revolves around changing reinforcement contingencies: one of an array of actions (e.g. pressing the left most lever) is paired with an outcome (e.g. receiving a food reward), and the subject learns to discriminate between those contingencies (Izquierdo & Jentsch, 2011). After reaching an accuracy criterion, the contingencies are reversed (e.g., only pressing the right lever, not the left, now leads to the food reward). Subjects must demonstrate cognitive and behavioral flexibility by constraining their previous responses and discarding the initially learned rule in order to maximize reward. Greater difficulty with stopping or updating behavior during reversal learning has been suggested to reflect greater impulsive action. Studies have shown that this behavioral inflexibility is genetically linked to impulsivity (Franken et al., 2008; Izquierdo & Jentsch, 2011). In addition to responding after reversal of response contingency, premature (inter-trial interval) responding within reversal learning can be measured. This measure is analogous to premature responding in the 5-CSRTT. Both measures of anticipatory responses are considered to be waiting impulsivity, or the inability to withhold response in anticipation of a reward-related cue (Dalley, Everitt, & Robbins, 2011).

Inbred mouse panels permit estimates of heritability of these traits, which is operationally defined as the proportion of phenotypic variation that is explained by genetic variation. In inbred lines, heritability is estimated as the proportion of phenotypic variance accounted for by strain. Past genetic reference population studies have considered this to be an effective estimate of heritability, considering each mouse from each strain is, to the extent maximally possible, genetically identical to one another (Hegmann & Possidente, 198; Philip et al., 2009). Environmental and technical sources of variance are reduced within these panels, further increasing the ability to detect heritability of a trait over external influence and providing an advantage over human twin studies (Williams et al., 2004).

Past efforts have utilized recombinant inbred (RI) panels to identify the heritability and genetic architecture of impulsive phenotypes. Laughin et al. (2011) conducted reversal learning in 51 BXD mouse strains and identified heritable strain variation in impulsivity, as well as a genome-wide significant quantitative trait locus (QTL) on chromosome 10. Positional candidate genes, including *Syn3* – encoding the synaptic phosphoprotein Synapsin III - expressed from this QTL were selected based upon expression phenotypes that were genetically correlated with the behavioral phenotype. A subsequent study using the BXD strains that exhibited extreme differences in reversal learning found that the poor reversal learning strains more rapidly acquired cocaine self-administration and administer cocaine at greater rates than do good reversal learning strains (Cervantes, Laughlin & Jentsch, 2013).

Utilizing mouse populations with greater genetic diversity may expand observed phenotypic ranges and lead to new insights in neurogenetics and neurobiology of impulsivity and its relationship to SUDs. The Collaborative Cross is a genetically diverse, multi-parental RI panel. It alleviates genetic bottleneck limitations by through the intercross of five classical inbred strains (A/J, C57BL/6J, 129S1/SvImJ, NOD/LtJ, NZO/HlLtJ), and three wild-derived inbred strains (CAST/EiJ, PWK/PhJ, WSB/EiJ), to capture >90% of the known genetic variation present in the laboratory mouse (Odet et al., 2015). In addition to having a large amount of genetic diversity, the Collaborative Cross also has generally balanced allele frequencies and evenly distributed recombination sites (Aylor et al., 2011; Philip et al., 2011), resulting in greater statistical power to detect genetic correlations among phenotypes gathered in different laboratories.

The present study aims to examine heritability and genetic correlations between locomotor response to novelty, palatable food consumption (a measure of reward sensitivity), reinforcement learning, two tests of impulsivity (delay discounting and reversal learning), and other catalogued addiction-related behaviors measured by others in the eight CC founder strains and ten CC strains. The tests of impulsivity are designed to measure three types of impulsivity: impulsive action (total trials to criteria in reversal learning), waiting impulsivity (anticipatory responses measured during reversal learning) and impulsive choice (indices of preference for an immediate reward). We hypothesized that the three measures of impulsivity are heritable traits, though not necessarily coinherited (genetically correlated). Locomotor activity and palatable food consumption are similarly anticipated to be heritable traits, each of which has shown conflicting evidence in the literature of being positively correlated with impulsivity (references?). These studies were conducted as part of a multi-lab collaboration in the Center for Systems Neurogenetics of Addiction. A set of strains were tested across different laboratories for impulsivity traits as well as SUD-related traits. Results from these efforts will provide insights into the inter-lab replicability of impulsivity traits as well as the genetic relationships among a large number of traits that may be predictive of SUDs.

### Subjects

All inbred and recombinant inbred mice involved in this study were born at The Jackson Laboratory (JAX; Bar Harbor ME). A subset was subsequently shipped to Binghamton University, while the remainder were phenotyped at JAX, in the facilities of the Behavioral Phenotyping Core of the Center for Systems Neurogenetics of Addiction. All procedures were performed according to the “Guide for the Care and Use of Laboratory Animals” (National Research Council, 2011) in the AAALAC accredited programs at Binghamton University or JAX, after approval by the relevant Institutional Animal Care and Use Committees.

### Binghamton University Site

Subjects were shipped from JAX to Binghamton University between 35-49 days of age. A total of 6 cohorts were shipped between 8/2016 and 8/2019. Mice from the eight founder strains were delivered in the first three cohorts, each of which was balanced to include representatives from all 8 strains and both sexes. Ten strains of CC mice were also studied; they were first included in cohort three. Details on mouse strains/cohorts are provided in Table 1.

**Table 1.** Strains tested at the Binghamton University site.

| **Mouse ID** | **JAX Stock No.** | **MGI ID** | **N** | **Cohorts** |
| --- | --- | --- | --- | --- |
| NOD/ShiLtJ | 001976 | 2162056 | 18 | 1-3 |
| 129S1/SvImJ | 002448 | 2160041 | 18 | 1-3 |
| A/J | 000646 | 2159747 | 18 | 1-3 |
| C57BL/6J | 000664 | 3028467 | 18 | 1-3 |
| NZO/HILtJ | 002105 | 2668669 | 18 | 1-3 |
| CAST/EiJ | 000928 | 2159793 | 18 | 1-3 |
| WSB/EiJ | 001145 | 2160667 | 18 | 1-3 |
| PWK/PhJ | 003715 | 2160654 | 18 | 1-3 |
| CC002/UncJ | 021236 | 5649080 | 12 | 4-6 |
| CC004/TauUncJ | 020944 | 5649082 | 23 | 3-6 |
| CC006/TauUncJ | 022869 | 5649237 | 12 | 4-6 |
| CC011/UncJ | 018854 | 5649240 | 12 | 4-6 |
| CC012/GeniUncJ | 028409 | 5694080 | 12 | 4-6 |
| CC025/GeniUncJ | 018857 | 5649246 | 11 | 4-6 |
| CC028/GeniUncJ | 025126 | 5659485 | 12 | 4-6 |
| CC032/GeniUncJ | 020946 | 5649248 | 12 | 4-6 |
| CC041/TauUncJ | 021893 | 5649251 | 24 | 3-6 |
| CC061/GeniUncJ | 023826 | 5649258 | 12 | 4-6 |

Upon arrival, mice were housed in the same groups in which they were shipped, with three mice of the same strain and sex being grouped together in a cage. The colony room was maintained on a 12h light/dark cycle (lights on at 0615 h) and at an average temperature of 69-70°F. During this initial acclimation period, food (Lab Diet 5001, ScottPharma Solutions) and water was available *ad libitum.* A nestlet and a translucent red acrylic tube (9.75cm long, 5cm diameter, approximately 65g) were placed in each cage. For removal from the cage, the mice were briefly handled by their tails using a gloved hand. Wild-derived inbred strains and some CC strains (CC004/TauUncJ and CC011/UncJ) were handled by the tail with forceps.

All three wild-derived strains (CAST/EiJ, WSB/EiJ, and PWK/PhJ) exhibit high levels of aggression and are at high risk of injury when group-housed in adulthood. To avoid injury and maintain uniformity across strains, all mice of both sexes and from all strains were singly housed in identical caging conditions at PND 60. Mice were acclimated to these conditions for 10 days until PND 70.

Prior to the initiation of the operant conditioning protocols described below, mice were introduced to a schedule of limited access to chow. Mice were weighed daily during food restriction and percent of free-feeding body weight was calculated by dividing the current weight by the free-feeding weight. During the limited access to food period, mice were fed once a day between 3pm and 5pm; chow quantity provided per day was titrated until mice reach 80-85% or 83-88% (CAST/EiJ, PWK/PhJ, WSB/EiJ) of their free feeding weights. Wild-derived inbred strains were maintained at a higher free-feeding percentage due to observed lethargy and dehydration when their weights approached 80%. Once mice reached their target weights, operant testing began (see Table 2 for reversal learning mouse weights; Table 3 for delay discounting mouse weights). If, at any point during the testing period, a mouse dropped below 80% of its free feeding weight, the quantity of chow provided was increased. If increased food availability did not lead to a recovery of body weight to ≥ 80% within a day, it was temporarily returned to *ad libitum* food access until its weight had recovered.

**Table 2.** Average of strain’s body weight during each stage of reversal learning testing, represented as a percentage of their initial free-feeding weight. SEM is represented as ± the mean.

| Mouse ID | Start | Training | Acquisition | Reversal |
| --- | --- | --- | --- | --- |
| NOD/ShiLtJ | 27.2 ± 0.99g | 85.8 ± 0.76% | 84.2 ± 0.71% | 85.8 ± 0.74% |
| 129S1/SvImJ | 23.7 ± 0.64g | 88.0 ± 1.04% | 86.6 ± 1.09% | 85.6 ± 1.02% |
| A/J | 21.8 ± 0.61g | 87.8 ± 0.62% | 84.7 ± 0.65% | 84.8 ± 0.52% |
| C57BL/6J | 23.6 ± 0.51g | 85.6 ± 0.47% | 84.7 ± 0.61% | 85.1 ± 0.54% |
| NZO/HiLtJ | 41.0 ± 1.24g | 88.4 ± 1.30% | 83.7 ± 0.92% | 83.6 ± 0.64% |
| CAST/EiJ | 15.5 ± 0.38g | 86.0 ± 0.92% | 85.3 ± 0.86% | 84.4 ± 0.65% |
| WSB/EiJ | 15.7 ± 0.40g | 86.7 ± 0.92% | 86.3 ± 0.77% | 87.6 ± 0.74% |
| PWK/PhJ | 17.2 ± 0.40g | 85.5 ± 0.43% | 85.7 ± 0.67% | 85.1 ± 0.61% |
| CC001/UncJ | 24.3 ± 0.90g | 84.1 ± 0.94% | 83.1 ± 0.88% | 82.8 ± 0.92% |
| CC002/UncJ | 24.2 ± 2.07g | 86.7 ± 0.66% | 85.5 ± 0.56% | 84.8 ± 0.86% |
| CC003//UncJ | 27.7 ± 0.59g | 84.0 ± 1.04% | 84.8 ± 0.98% | 84.9 ± 1.15% |
| CC004/TauUncJ | 28.6 ± 1.44g | 86.8 ± 1.76% | 84.7 ± 0.94% | 85.6 ± 1.45% |
| CC005/TauUncJ | 19.9 ± 0.92g | 87.1 ± 0.89% | 85.8 ± 0.86% | 85.9 ± 0.57% |
| CC006/TauUncJ | 23.8 ± 0.98g | 86.0 ± 1.04% | 87.0 ± 0.71% | 86.9 ± 0.57% |
| CC008/GeniUncJ | 32.4 ± 0.73g | 84.7 ± 1.01% | 86.5 ± 0.86% | 85.7 ± 0.85% |
| CC011/UncJ | 25.7 ± 1.01g | 87.6 ± 0.88% | 85.5 ± 1.09% | 85.4 ± 0.73% |
| CC012/GeniUncJ | 20.8 ± 1.38g | 87.3 ± 1.21% | 86.6 ± 1.17% | 87.9 ± 0.49% |
| CC013/GeniUncJ | 27.2 ± 1.01g | 83.8 ± 0.88% | 86.0 ± 0.98% | 86.8 ± 0.77% |
| CC015/UncJ | 21.0 ± 1.36g | 86.1 ± 0.78% | 83.5 ± 0.77% | 84.8 ± 0.90% |
| CC016/GeniUncJ | 24.6 ± 1.87g | 85.0 ± 0.33% | 83.2 ± 0.49% | 82.7 ± 0.36% |
| CC017/UncJ | 21.0 ± .59g | 85.3 ± 0.67% | 83.9 ± 0.93% | 84.4 ± 0.90% |
| CC019/TauUncJ | 19.1 ± 2.91g | 86.3 ± 0.27% | 84.7 ± 2.25% | 83.8 ± 2.17% |
| CC023/GeniUncJ | 22.5 ± 1.23g | 84.4 ± 85.1% | 85.1 ± 2.48% | 82.2 ± 1.59% |
| CC025/GeniUncJ | 23.6 ± 1.43g | 86.1 ± 0.32% | 83.1 ± 1.83% | 83.1 ± 0.50% |
| CC026/GeniUncJ | 25.3 ± 1.79g | 84.8 ± 0.50% | 84.5 ± 0.83% | 82.9 ± 0.65% |
| CC027/GeniUncJ | 24.4 ± 1.40g | 86.9 ± 0.50% | 85.1 ± 1.11% | 83.7 ± 1.13% |
| CC028/GeniUncJ | 29.4 ± 2.03g | 86.3 ± 0.57% | 84.5 ± 0.27% | 83.8 ± 1.18% |
| CC030/GeniUncJ | 23.9 ± 1.23g | 86.4 ± 0.50% | 85.2 ± 0.20% | 85.7 ± 0.44% |
| CC032/GeniUncJ | 25.7 ± 1.60g | 90.7 ± 1.55% | 86.0 ± 0.75% | 86.0 ± 0.00% |
| CC033/GeniUncJ | 21.7 ± 1.77g | 82.8 ± 0.59% | 85.4 ± 1.27% | 84.7 ± 0.86% |
| CC036/UncJ | 25.7 ± 1.75g | 85.4 ± 0.66% | 85.2 ± 1.87% | 84.5 ± 1.57% |
| CC037/TauUncJ | 22.8 ± 0.80g | 84.7 ± 2.15% | 84.8 ± 0.15% | 85.3 ± 0.90% |
| CC040/TauUncJ | 32.9 ± 3.60g | 86.1 ± 0.10% | 81.7 ± 1.35% | 79.4 ± 0.25% |
| CC041/TauUncJ | 24.9 ± 1.42g | 88.1 ± 0.84% | 85.3 ± 0.92% | 85.8 ± 0.13% |
| CC043/GeniUncJ | 20.1 ± 0.00g | 84.5 ± 1.05% | 84.4 ± 2.85% | 86.1 ± 2.45% |
| CC051/TauUncJ | 32.9 ± 3.27g | 85.2 ± 0.49% | 83.5 ± 0.88% | 82.9 ± 0.32% |
| CC057/UncJ | 28.3 ± 2.25g | 83.0 ± 0.70% | 82.4 ± 0.05% | 84.0 ± 0.30% |
| CC059/TauUncJ | 35.1 ± 2.60g | 83.9 ± 1.00% | 82.0 ± 0.15% | 82.6 ± 0.50% |
| CC061/GeniUncJ | 21.8 ± 0.00g | 87.4 ± 0.00% | 85.1 ± 0.00% | 87.2 ± 0.00% |
| CC075/UncJ | 25.9 ± 1.70g | 85.8 ± 0.15% | 81.4 ± 0.20% | 81.7 ± 0.20% |
| CC078/TauUncJ | 25.7 ± 2.05g | 87.7 ± 1.25% | 87.7 ± 3.00% | 86.1 ± 1.25% |
| CC079/TauUncJ | 21.2 ± 3.70g | 85.3 ± 0.60% | 83.8 ± 0.75% | 84.2 ± 0.40% |
| CC080/TauUncJ | 20.1 ± 5.05g | 84.0 ± 1.05% | 86.5 ± 0.05% | 84.0 ± 0.10% |
| CC084/TauJ | 28.7 ± 2.55g | 83.7 ± 0.15% | 83.0 ± 0.65% | 83.7 ± 0.60% |

**Table 3.** Average of strain’s body weight during each stage of delay discounting testing, represented as a percentage of their initial starting weight (grams). No significant differences between strains were observed. SEM is represented as ± the mean.

| Mouse ID | Start | 0s | 3s | 6s | 9s |
| --- | --- | --- | --- | --- | --- |
| NOD/ShiLtJ | 27.4 ± 1.98g | 89.0 ± 1.13% | 89.3 ± 1.10% | 88.7 ± 0.75% | 88.9 ± 1.07% |
| 129S1/SvImJ | 22.5 ± 0.78g | 89.8 ± 1.42% | 87.7 ± 1.17% | 90.1 ± 1.39% | 89.8 ± 1.61% |
| A/J | 21.7 ± 0.54g | 88.9 ± 1.32% | 90.8 ± 1.19% | 91.3 ± 0.61% | 91.0 ± 0.96% |
| C57BL/6J | 24.2 ± 1.20g | 90.2 ± 1.23% | 85.5 ± 0.86% | 89.5 ± 1.32% | 88.7 ± 0.95% |
| NZO/H1LtJ | 39.7 ± 1.15g | 88.6 ± 1.07% | 89.2 ± 1.16% | 89.8 ± 0.89% | 87.8 ± 0.95% |
| CAST/EiJ | 15.5 ± 0.56g | 90.0 ± 1.32% | 89.8 ± 0.77% | 91.5 ± 1.06% | 90.5 ± 0.88% |
| WSB/EiJ | 17.2 ± 0.58g | 89.1 ± 1.15% | 89.3 ± 1.13% | 88.2 ± 1.04% | 90.3 ± 1.35% |
| PWK/PhJ | 16.0 ± 0.57g | 91.4 ± 0.94% | 86.8 ± 0.75% | 89.7 ± 0.79% | 88.2 ± 1.31% |
| CC002/UncJ | 26.0 ± 0.79g | 87.8 ± 0.85% | 90.5 ± 1.25% | 88.2 ± 1.26% | 88.5 ± 0.93% |
| CC004/TauUncJ | 26.9 ± 1.14g | 87.8 ± 0.96% | 87.6 ± 1.03% | 88.3 ± 1.26% | 87.8 ± 1.23% |
| CC006/TauUncJ | 24.0 ± 0.77g | 88.7 ± 1.65% | 90.7 ± 1.51% | 90.0 ± 0.76% | 87.0 ± 1.41% |
| CC011/UncJ | 27.7 ± 1.55g | 88.7 ± 1.05% | 89.7 ± 1.88% | 88.5 ± 1.87% | 88.5 ± 1.83% |
| CC012/GeniUncJ | 21.0 ± 1.05g | 90.5 ± 1.19% | 89.0 ± 1.75% | 91.5 ± 0.98% | 87.5 ± 1.38% |
| CC025/GeniUncJ | 25.1 ± 1.13g | 88.4 ± 1.66% | 87.0 ± 2.17% | 92.4 ± 1.63% | 88.6 ± 1.44% |
| CC028/GeniUncJ | 27.5 ± 1.20g | 89.7 ± 1.33% | 88.0 ± 1.82% | 89.7 ± 2.35% | 85.3 ± 0.91% |
| CC032/GeniUncJ | 21.0 ± 0.39g | 92.3 ± 0.99% | 88.0 ± 2.01% | 88.8 ± 1.29% | 86.8 ± 1.37% |
| CC041/TauUncJ | 25.9 ± 1.09g | 86.4 ± 2.01% | 89.1 ± 1.37% | 87.5 ± 0.95% | 89.7 ± 1.09% |
| CC061/GeniUncJ | 21.2 ± 0.93g | 92.2 ± 1.69% | 87.8 ± 1.05% | 88.5 ± 1.34% | 90.2 ± 1.54% |

### The JAX Site

With the following exceptions, maintenance and feeding of the mice at JAX followed the same details outlined above. The eight CC founder strains were bred and maintained within the Behavioral Phenotyping Core of the CSNA at JAX; all CC strains were born in the commercial Production facility at JAX and were relocated to the Behavioral Phenotyping Core at four to six weeks of age. Mouse home cages contained a disposable dome-shaped shack (Shepherd Specialty Papers, Inc., Watertown, TN, USA), instead of a red tube. A total of eight founder strains and thirty-six CC strains were studied (Table 4), with thirty-six CC strains tested at the JAX site, and ten of those strains tested simultaneously at the Binghamton site.

**Table 4.** Strains tested at the Jackson Laboratory site.

| **Mouse ID** | **JAX Stock No.** | **MGI ID** | **N** | **Source** |
| --- | --- | --- | --- | --- |
| NOD/ShiLtJ | 001976 | 2162056 | 18 | Bred in-house |
| 129S1/SvImJ | 002448 | 2160041 | 7 | Bred in-house |
| A/J | 000646 | 2159747 | 10 | Bred in-house |
| C57BL/6J | 000664 | 3028467 | 27 | Bred in-house |
| NZO/HiLtJ | 002105 | 2668669 | 7 | Bred in-house |
| CAST/EiJ | 000928 | 2159793 | 19 | Bred in-house |
| WSB/EiJ | 001145 | 2160667 | 15 | Bred in-house |
| PWK/PhJ | 003715 | 2160654 | 17 | Bred in-house |
| CC001/UncJ | 021238 | 5649079 | 4 | Shipped |
| CC002/UncJ | 021236 | 5649080 | 6 | Shipped |
| CC003//UncJ | 021237 | 5649081 | 4 | Shipped |
| CC004/TauUncJ | 020944 | 5649082 | 3 | Shipped |
| CC005/TauUncJ | 020945 | 5649083 | 4 | Shipped |
| CC006/TauUncJ | 022869 | 5649237 | 4 | Shipped |
| CC008/GeniUncJ | 026971 | 5659476 | 3 | Shipped |
| CC011/UncJ | 018854 | 5649240 | 6 | Shipped |
| CC012/GeniUncJ | 028409 | 5694080 | 6 | Shipped |
| CC013/GeniUncJ | 021892 | 5649241 | 4 | Shipped |
| CC015/UncJ | 018859 | 5659478 | 2 | Shipped |
| CC016/GeniUncJ | 021237 | 5659479 | 8 | Shipped |
| CC017/UncJ | 022870 | 5649242 | 4 | Shipped |
| CC019/TauUncJ | 021894 | 5649244 | 10 | Shipped |
| CC023/GeniUncJ | 025131 | 5659483 | 1 | Shipped |
| CC025/GeniUncJ | 018857 | 5649246 | 8 | Shipped |
| CC026/GeniUncJ | 024685 | 5649247 | 2 | Shipped |
| CC027/GeniUncJ | 025130 | 5659484 | 5 | Shipped |
| CC028/GeniUncJ | 025126 | 5659485 | 7 | Shipped |
| CC030/GeniUncJ | 025426 | 5659487 | 7 | Shipped |
| CC032/GeniUncJ | 020946 | 5649248 | 2 | Shipped |
| CC033/GeniUncJ | 025910 | 5659488 | 2 | Shipped |
| CC036/UncJ | 025127 | 5659490 | 6 | Shipped |
| CC037/TauUncJ | 025423 | 5649249 | 3 | Shipped |
| CC040/TauUncJ | 023831 | 5649250 | 3 | Shipped |
| CC041/TauUncJ | 021893 | 5649251 | 3 | Shipped |
| CC043/GeniUncJ | 023828 | 5649256 | 2 | Shipped |
| CC051/TauUncJ | 021897 | 5649257 | 2 | Shipped |
| CC057/UncJ | 024683 | 5659646 | 2 | Shipped |
| CC059/TauUncJ | 025125 | 5659650 | 10 | Shipped |
| CC061/GeniUncJ | 023826 | 5649258 | 6 | Shipped |
| CC075/UncJ | 027293 | 5659668 | 3 | Shipped |
| CC078/TauUncJ | 025989 | 5796475 | 2 | Shipped |
| CC079/TauUncJ | 025990 | 5796476 | 2 | Shipped |
| CC080/TauUncJ | 025988 | 5796471 | 3 | Shipped |
| CC084/TauJ | 028923 | 6159255 | 3 | Shipped |

### Phenotyping Protocols

The Binghamton University cohort was sequentially phenotyped using a set of protocols described below. They were evaluated first for <u>Locomotor Response to Novelty and Habituation</u> (PND 70-71), followed by testing for <u>Palatable Food Consumption</u> (PND 72-78) and then one of two operant conditioning procedures: either <u>Reversal Learning</u> or <u>Delay Discounting</u>. The cohort evaluated at JAX were sequentially phenotyped in open field, light/dark, hole board, and novelty place preference assays prior to Reversal Learning. At both sites, behavioral tests were conducted during the light phase of the animal’s light cycle.

### Locomotor Response to Novelty and Habituation (Binghamton University pipeline)

At PND 70, mice were assessed for locomotor response in a novel open field environment. Subjects were transported to a testing room on a cart. Each mouse was individually placed in a 17 “L × 17” W × 12“ H (43.2 × 43.2 × 30.5 cm) open field chamber fitted with infrared beams (Med-Associates MED-OFAS-RSU; St Albans VT). All open field chambers were within sound attenuating cubicles measuring 26” W × 22“ H × 20.5” D (66 × 52.7 × 55.9 cm) at the interior with walls 0.75“ (1.9 cm) thick. Activity was recorded for 40-min. The primary dependent measure examined was total distance traveled in cm, as well as the duration of time (in s) spent: ambulating, resting, performing stereotypy, or rearing. After the session, mice were immediately removed and returned to their home cage. Between subjects, the apparati were cleaned with a mixture of 10% Alconox detergent in water.

The following day at the same time, mice were placed back in the same open field chamber, and activity parameters were again recorded for 40-min. Locomotor habituation was assessed by calculating the difference in total distance traveled between day 1 and 2.

#### Palatable Food Consumption

Beginning on PND 72 (one day after the conclusion of locomotor assessments), mice experienced access to a highly palatable chocolate-flavored Boost solution (Nestle) in their home cage. Boost was made available using a plastic petri dish that was placed on top of the bedding. The solution was available continuously for a 48-h period, with the solution being refreshed at the 24-h time point.

From PND 74 to 80, mice were evaluated daily for Boost (and water) consumption in 2-h lickometry sessions. All testing took place inside dual lickometer Scurry boxes (Model 80822S, activity wheel removed; Lafayette Instruments, Lafayette IN). Each lickometer box is 35.3 × 23.5 × 20c m and is fitted with a food hopper and two 50 mL sipper bottles, with pine chip bedding on the chamber floor. Mice were transported to the testing room on a cart. Room lights were on during testing and a room dehumidifier provided ambient background noise. Mice were placed, individually, into the lickometer boxes and allowed to freely consume Boost and water (no chow provided). One bottle was filled with Boost solution and the other was filled with water; the position (left or right) of the two solutions relative to one another was counterbalanced pseudorandomly across the testing days. Licks on each spout were counted by a computer. The number of licks was divided by the body weight of the animal to account for behavioral variability that might be attributable to body weight differences.

#### Operant Conditioning – Reversal Learning

Following Boost consumption testing, mice were maintained on food-restriction and transitioned to operant testing when a stable target weight was achieved. Half of the mice from each sex and each strain were randomly designated for evaluation using an operant discrimination/reversal learning procedure. All operant testing took place in 8.5” L × 7” W × 5” H (21.6 × 17.8 × 12.7 cm) operant modular chambers (Model ENV-307W, Med Associates Inc.) that were fitted with stainless-steel grid floors (Model ENV-307W-GFW, Med Associates Inc.) and located in sound attenuating cubicles. The operant box contains a horizontal array of five nose poke apertures on one side of the box, and a central food magazine, flanked on the left and right by two retractable, ultrasensitive levers on the other. A house light and white noise maker are positioned within the cubicle above the operant box. Mice were removed from their home cage by their tail and placed inside the operant box. To remove the CAST/EiJ, PWK/PhJ, and WSB/EiJ wild-derived strains and the CC004/TauUncJ and CC011/UncJ from the operant chamber, the red tube from their home cage was placed into the box; once the mouse entered the tube, it was transferred into their home cage.

Each mouse was sequentially tested in a series of programs; mice transitioned from program to program individually, as they met criterion performance (see below). Mice underwent the following programs:

Stage 1: **Box habituation**. House light and white noise were active. No reinforcements were provided. Box habituation comprised of one session that lasted 1-h.

Stage 2: **Magazine training**. House light and white noise were active for the duration of the test. During this test, 20-21µl Boost was dispensed into the food magazine every 30 s. The session ended after 1-h or after the mouse received and retrieved 50 rewards, whichever came first. A mouse progressed to Stage 3 when it earned 30 or more rewards within a session.

Stage 3: **Initial operant (nose-poke) conditioning**. Sessions began with illumination of the house light and activation of the white noise generator; 10-s later, nose poke aperture 3 of 5 (center aperture) was illuminated. A behavioral response that broke the photocell in the aperture (usually, a nose poke) resulted in the extinction of the internal light; in addition, if the beam was broken for a continuous pre-set period of at least 0 (beam break with no additional hold time), 100, or 200 ms (the time requirement varied randomly from trial to trial), the action was reinforced by the delivery of 20-21µl of Boost solution; after each reinforcer was retrieved, a new trial was initiated 1.5-s later (signaled by illumination of the center nose poke aperture). If a response was initiated but was not sustained for the preset period, a time out period of 2-s occurred, during which time the central nose poke light and house light were extinguished. If a mouse did not voluntarily respond in the center hole for at least 15 minutes, the center hole was baited with a Boost-saturated cotton swab. Daily sessions lasted up to 1-h but were also terminated if an individual mouse earned 50 reinforcers. Each mouse was tested daily on this stage until it received at least 50 reinforcers in a single session, at which time it progressed to the next stage.

Stage 4: Mice were tested under the same basic conditions outlined in Stage 3, except that a minimum duration nose poke of 100- or 200-ms was required to produce reinforcement. If a mouse had not responded in the central illuminated hole for 15 minutes, the center hole was again baited with a Boost-saturated cotton swab. When the mouse earned 50 reinforcers in a single session, it progressed to Stage 5. If the mouse had not met criteria after 10 days, it was regressed to Stage 3. If the subject returned to Stage 4 but still did not meet criteria after another 10 test days, they were removed from the study due to failure to progress. Across all Stages, a mouse could only regress once. For example, if a mouse did not pass Stage 4 in 10 days and regressed to Stage 3, then later did not pass Stage 5 within 10 days, the mouse was removed from the study.

Stage 5: In this phase, mice were tested under the same basic conditions as outlined in Stages 3 and 4, except that a minimum duration nose poke of 100-, 200-, or 300-ms was required to trigger reinforcement delivery. If a mouse did not respond in the center illuminated hole for 15 minutes, the center hole was baited with a Boost-saturated cotton swab. When the mouse earned 50 reinforcers in a single session, it progressed to the **Discrimination Learning** stage. If the mouse did not meet passing criteria after 10 days, they regressed to Stage 4. If the subject returned to Stage 5 but still did not meet criteria after another second of 10 test days, they were removed from the study due to regression failure.

Stage 6: **Discrimination learning**. As above, session onset was signaled by illumination of the house light and activation of the white noise generator; trial onset was signaled by illumination of the center nose poke aperture. As in Stage 5, mice were required to first complete an observing response into the central aperture of 100-, or 200-ms duration; any nosepokes into the target (flanking) holes before completing the observing response and successfully initiating a trial were counted as premature/anticipatory responses. Once a trial was successfully initiated with an observing response, the two apertures flanking the central hole (hole 2 and 4) were immediately illuminated. A response into one of the two apertures (pseudorandomly assigned across strains) resulted in the delivery of a Boost reinforcer (this was counted as a correct choice). Poking into the other hole - or not making any response within 30-s, triggered a time out, during which time the house light was extinguished; these outcomes were counted as an incorrect choice or an omission, respectively. Daily sessions of 1-hr were conducted until learning criteria were met; these criteria included a mouse completing at least 20 trials in a single session and achieving at least 80% accuracy over a running window including the last 20 trials. A mouse is regressed to Stage 5 if it does not complete at least 10 trials for three consecutive days. If 300 trials are conducted without meeting passing criteria, the mouse is removed from the study due to Stage 6 failure.

Stage 7: **Reversal learning stage**. Testing was nearly identical to that described above in Stage 6, with the exception that the reinforcement contingencies associated with the two holes were switched. Testing progressed in daily sessions until animals once again met the same learning criteria rule described above, and the same dependent variables were collected (see below). After reversal was completed, mice were gradually adjusted back onto an *ad libitum* feeding schedule. Subjects were removed from the study due to Stage 7 failure if 400 trials were conducted without meeting criteria or if 8 weeks of testing passed.

Key dependent variables for the discrimination learning and reversal learning stages were: total trials required to reach criteria (TTC) in each stage and premature responding. TTC was calculated as the total number of completed trials (all trials ending in an incorrect or correct response) until it met the performance criteria. The difference in TTC in the reversal stage to TTC in the discrimination learning stage demonstrates each animal’s ability to alter responding under a changing reward contingency, with a non-zero, positive difference score indicating difficulty with altering its behavior and/or inhibiting the initially trained response.

Premature responses are nose pokes into one of the flanking target holes before a trial is successfully initiated, a measure roughly analogous to that collected in the 5-choice serial reaction time (Bari et al. 2008). Premature responses were separately counted for the correct and incorrect target aperture. Premature responding thus has four values: premature responding in the correct hole at acquisition, premature responding in the incorrect hole at acquisition, premature responding in the correct hole at reversal, and premature responding in the incorrect hole at reversal. All premature responding values were further divided by the animal’s TTC in that stage to estimate the average number of premature responses made per trial. Of particular interest is premature responding in the correct hole during acquisition and in the incorrect hole at reversal, as these are the dominant types of responses made.

Other variables measured were the frequency of omissions (total omissions/TTC for each stage); average correct trials (total correct trials/TTC; average trial initiation latency (total trial initiation latency/TTC), which is the average amount of time that passes between the end of one trial and the successful initiation of the next one; and average reward retrieval time (total reward retrieval time/total correct trials), which is the average amount of time that passes between a reward being administered and the animal’s head entering the magazine.

#### Operant Conditioning – Delay Discounting

As described above, mice were transitioned to a limited food access schedule, once lickometer testing was completed. Once targeted reductions in body weights were achieved, half of the mice from each sex and strain were randomly designated for evaluation using a delay discounting procedure.

Each mouse was sequentially tested in a series of training stages through which they transitioned individually, as they met criterion performance (see below). Mice underwent the following programs:

Stage 1: **Box habituation**. House light and white noise were active. No reinforcements were provided. Box habituation comprised of one session that lasted 1-h.

Stage 2: **Magazine training**. Again, the house light and white noise were active for the duration of the test, and 20-21µl Boost is dispensed every 30 s. The duration of testing was 1 h. A mouse progressed to Stage 3 when it earned 30 or more rewards within a session.

Stage 3: **Initial operant (lever press) conditioning**. Session onset was signaled by illumination of the house light and activation of the white noise generator. On each trial, one lever (left or right) was inserted to the chamber and actuation of the lever by the mouse triggered delivery of 20-21 µl of Boost on a fixed-ratio 1 schedule of reinforcement. Across trials, the lever that was inserted (left or right) was pseudorandomly varied, such that each mouse actuated each lever a roughly equal number of times. Each daily session ended after 1-h or after 60 reinforcers were obtained, whichever came first. A mouse progressed to Stage 4 when it earned 60 reinforcers within a session.

Stage 4: In this stage of lever press training, a procedure nearly identical to that used in Stage 3, described above, was employed. The only difference was that responses on the inserted lever were reinforced on a fixed ratio 3 schedule of reinforcement. The program ended after 1-h or after 60 rewards were obtained. A mouse progressed to Stage 5 when it earned 60 reinforcers within a session.

Stage 5: **Observing response training**. The only difference from Stage 4 was that the mouse was required to complete an observing response (nose poke response into the central nose-poke hole [aperture 3 of 5] on the side of the chamber opposite to the levers in order to trigger insertion of a lever). Responses on that lever were still reinforced on a fixed-ratio 3 schedule. The program ended after 1-h or after 60 rewards were obtained. A mouse progressed to Stage 6 when it earned 60 reinforcers within a session.

Stage 6: In this stage of training, a procedure nearly identical to that described in Stage 5 was used. The only difference was that the program ended after 1.5 hours, or after 80 rewards were obtained, whichever came first. A mouse progressed to Side bias determination when it earned 80 reinforcers within a session.

Stage 7: **Side bias determination**. Trials began with presentation of both levers; responding on either lever was reinforced by 20-21 µl of Boost on a fixed-ratio 3 schedule. After a 10-s inter-trial interval, both levers are again presented, but only responses on the alternate lever were rewarded. A trial was only counted if the mouse successfully pressed the alternate lever. The program ended after 40 trials, or after 1.5 hours. The lever (right or left) on which each trial was initiated was recorded. The fraction of trials initiated with left versus right responses was determined, and the lever with the highest fraction was selected to be the one associated with the large-delayed outcome during the subsequent delay discounting testing. After a single session, a mouse progressed to Delay discounting assessment.

Stage 8: **Delay discounting assessment**. These daily tests began with activation of the house light and white noise generator. Individual trials commenced with activation of the light in aperture 3 of 5. A nose poke into the aperture triggered presentation of both levers. Completion of the FR3 schedule on one lever led to a small (10µl) reward delivered immediately (termed the SI option), while completion of the schedule on the other led to a larger (20µl) reward delayed by 0-9 s (termed the LD option). Delays were varied between session; all mice encountered delays of 0, 3, 6 or 9 s. Each delay was encountered for three consecutive days, and the order of delays experienced was controlled by a cycle Latin square design, balanced across strains. Reward magnitude was adjusted within session. Specifically, the reward volume delivered increased or decreased by 10% depending on the subject’s prior lever choice, such that choosing the SI lever would cause the SI reward to decrease by 10% and the LD to increase by 10% on the next trial, and vice versa if the LD lever is selected. Selecting the same reward lever five times in a row resulted in a forced choice trial, in which only the unchosen lever was ejected, forcing the subject to select that option. Reward amounts were not titrated for the forced trial and forced trials were not included in data analyses.

The final volume delivered on the SI lever was measured and subtracted from the final volume delivered on the LD lever; positive values indicate greater discounting, as the volume of the LD outcome needed to be substantially greater than that of the SI option, due to the delay. The reward magnitude for each option was averaged over the last 30 trials, and those values were averaged across the three days that delay was tested.

Once the first Latin square is completed, mice receive a two-day break, and they then experienced a second Latin square design varying delay across sessions.

### Data Analyses

Data for each variable was analyzed using SPSS Statistics (Version 23). Data was first examined using a body plot, and outliers two standard deviations from the mean within strain were removed. Statistical significance was established as a probability level of *p*<.05. Independent variables were strain and sex. A total of eighteen strains (eight founders and ten CC) was utilized for all analyses except for reversal learning, which included the eight founders and thirty-six CC strains.

Heritability estimates were derived for each significant main effect of strain using effect size, which is an estimate of the variance accounted for by the independent variable divided by the total amount of variance. The proportion of variance explained by strain reveals the portion of all phenotypic variation that is heritable (Philip et al., 2009).

*h^2^ = σ^2^*Between Strain / (*σ^2^*Within Strain + *σ^2^*Between Strain)

Strain-level correlations were conducted only on subjects (8 founder strains and 10 CC strains) at the Binghamton site. A Pearson’s correlation and Spearman’s correlation were used to analyze the relationship between key variables of interest from each behavioral test within this study, as well as key variables from within the study to other phenotypes characterized in these strains. Alpha level was adjusted with a Bonferroni correction to account for multiple comparisons.

## Results

Strains exhibited heritable variation in free-feeding body weights (Tables 2 and 3); ANOVA revealed an interaction of strain by sex (F[17,310]=1.720, *p*<.05), along with main effects of strain (F[17,310]=93.421, *p*<.001) and sex (F[1,310]=152.463, *p*<.001). Cohort effects were examined for Binghamton site mice and no effect was detected for TTC, premature responding in acquisition or reversal, or the 0s, 3s, and 6s delay discounting measures (*p*>.05 for all). A cohort effect was found for the 9s delay (F[1,5]=2.628, *p*<.05), driven by Cohort 2 exhibiting significantly lower 9s difference scores than the other cohorts.

### Locomotor Response to Novelty and Habituation

Total distance traveled in the open field chambers was assessed in 2 consecutive daily test sessions (Figure 1). Repeated measures ANOVA revealed an interaction between day and strain (F[17,24]= 5.306, *p*<.001, ηp^2^=.268) and a trending interaction of strain and sex (F[17,247]=1.627, *p*=.058, ηp^2^=.101); there were no other significant interactions involving strain. Main effects of day (F[1,247]=57.560, *p*<.001, ηp^2^=.189), strain (F[17,247]=9.828, *p*<.001, ηp^2^=.404) and sex (F[1,247]=8.015, *p*<.01, ηp^2^=.031) were also observed. Notably in the founder strains, two patterns of response were observed: strains with high ambulatory distance traveled (C57BL/6J, NOD/ShiLtJ, CAST/EiJ, PWK/PhJ, WSB/EiJ) and strains with low ambulatory distance traveled (NZO/HlLtJ, 129S1/SvImJ, and A/J). CC strains, on the other hand, demonstrated more continuous variation in ambulatory distance, a trait previously observed for behavioral wildness in this panel (Philip et al., 2011). Further analysis of the founder strains utilizing pairwise comparisons revealed that the CAST/EiJ, C57BL/6J, NZO/HlLtJ, and WSB/EiJ strains exhibited clear evidence of habituation with significantly less distance traveled on day 2 than on day 1 (all p<0.05), while the 129S1/SvImJ, A/J, PWK/PhJ, and NOD/ShiLtJ strains did not habituate.

**Figure 1.**
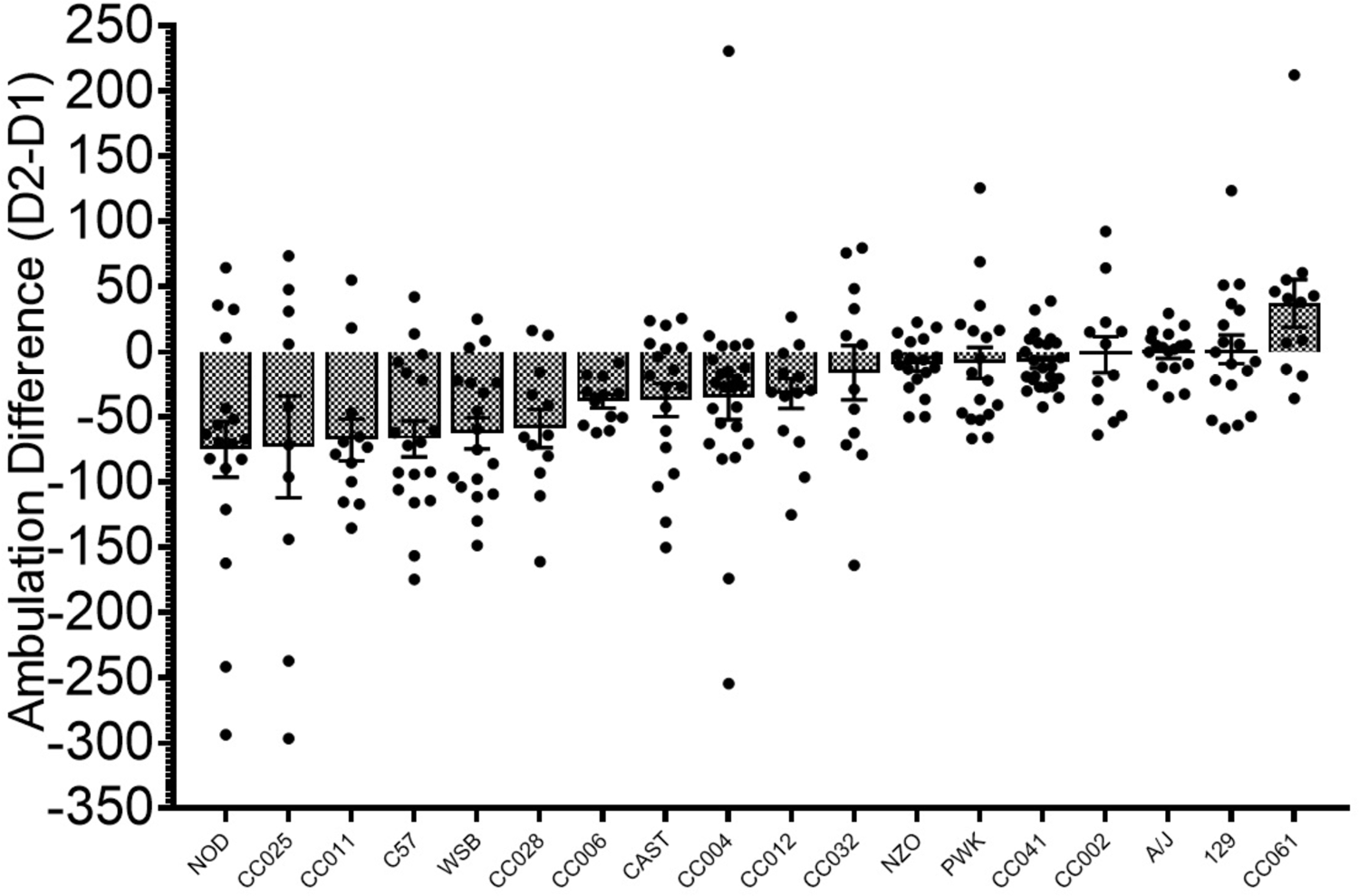
Ambulatory distance difference (Day 2-Day 1) calculated by total centimeters in the open field chamber.

Duration of time (s) spent ambulating, resting, performing stereotypy, or rearing was also examined. Results of these analyses (see Table 5) demonstrated consistent main effects of day and strain (with heritability estimates ranging from 0.452 to 0.695) for all measures. The effect of day again reveals habituation to novelty, in this case characterized by decreased ambulation and stereotypy and increased rest time and ambulation speed (Table 5). Strain significantly moderated observed habituation in ambulation, stereotypy, and rearing time, as evidenced by a day by strain interaction. Habituation of ambulation time was observed only in the CAST/EiJ, C57BL/6J, NOD/ShiLtJ, and WSB/EiJ strains; habituation of stereotypy was found in the A/J, CAST/EiJ, C57BL/6J, NOD/ShiLtJ, PWK/PhJ, and WSB/EiJ strains; and habituation of rearing was observed in the A/J, NOD/ShiLtJ, and NZO/HlLtJ strains.

**Table 5.** ANOVA statistics for duration of ambulation, resting, stereotypy and rearing (s), as well as ambulation speed (cm/s) during the open field test (40 min). Significant findings are bolded and italicized.

|  | Strain | Sex | Day | Strain x Sex | Strain x Day | Strain x Sex x Day |
| --- | --- | --- | --- | --- | --- | --- |
| Ambulation time | *F[17,250]=12.138, p<.001, $\eta p^2=.452$* | *F[1,250]=8.696, p<.01, $\eta p^2=.034$* | *F[1,250]=76.119, p<.001, $\eta p^2=.233$* | *F[17,250]=1.111, p<.01, $\eta p^2=.121$* | *F[17,250]=4.474, p<.001, $\eta p^2=.233$* | F[17,250]=1.244, p>.05 |
| Resting time | *F[17,250]=33.524, p<.001, $\eta p^2=.695$* | *F[1,250]=15.664, p<.001, $\eta p^2=.059$* | *F[1,250]=28.780, p<.001, $\eta p^2=.103$* | *F[17,250]=2.050, p<.01, $\eta p^2=.122$* | F[17,250]=1.262, p>.05 | F[17,250]=1.006, p>.05 |
| Stereotypy time | *F[17,250]=24.593, p<.001, $\eta p^2=.626$* | F[1,250]=2.230, p>.05 | *F[1,250]=2.050, p<.01, $\eta p^2=.122$* | F[17,250]=1.294, p>.05 | *F[17,250]=1.2346, p<.01, $\eta p^2=.138$* | *F[17,250]=1.697, p<.05, $\eta p^2=.103$* |
| Rearing time | *F[17,250]=28.083, p<.001, $\eta p^2=.656$* | *F[1,250]=50.445, p<.001, $\eta p^2=.168$* | *F[1,250]=17.884, p<.001, $\eta p^2=.067$* | *F[17,250]=2.589, p<.001, $\eta p^2=.150$* | *F[17,250]=4.690, p<.001, $\eta p^2=.242$* | *F[17,250]=1.824, p<.05, $\eta p^2=.110$* |
| Speed | *F[17,250]=17.102, p<.001, $\eta p^2=.538$* | F[1,250]=.012, p>.05 | *F[1,250]=5.182, p<.05, $\eta p^2=.02$* | F[1,250]=.976, p>.05 | F[17,250]=1.341, p>.05 | F[17,250]=.565, p>.05 |

Ambulation, resting, and rearing time phenotypes also revealed interactions between strain and sex (Table 5). Overall, the data reveals a trend for males and females to exhibit similar ambulatory phenotypes except in select strains. This is demonstrated by the PWK/PhJ and CC004/TauUncJ females that spent more time ambulating than did males from these strains; CAST/EiJ, WSB/EiJ, CC004/TauUncJ, and CC006/UncJ females spent less time resting than males; and NOD/ShiLtJ, C57BL/6J, CAST/EiJ, PWK/PhJ, CC004/TauUncJ, CC006/UncJ, CC061/GeniUncJ females spent less time rearing than males. Notably, the strains that show significant sex differences in exploration include all three of the wild-derived founder strains (CAST/EiJ, WSB/EiJ, and PWK/PhJ).

### Palatable Food Consumption

Spout contacts were analyzed within daily, 2-h sessions across 7 consecutive days of testing as well as averaged across the last three days of testing (days 5-7). Unadjusted lick data were analyzed as well as lick data adjusted for the number of licks by each animal’s body weight (Licks/Body weight) to control for the possibility that body size could influence strain difference in consummatory behavior. These adjusted and unadjusted values were compared with a Spearman’s correlation, and a significant positive correlation was detected (*r_s_*[16]=.872, *p*<.001), showing that the rank order of strain differences was not significantly altered by adjusting for body weight; consequently, values adjusted for body weight were used as the primary variable for analyses.

A repeated measures ANOVA on licks across the seven days identified main effects of day (F[6,228]=17.444, *p*<.001, ηp^2^=.071) and strain (F[17,228]=5.788, *p*<.001, ηp^2^=.541; Figure 2), and a trending interaction of day by strain (F[6,228]=1.232*, p*=.081, ηp^2^=.084). No effect of sex or interactions of sex with day or strain were found. Water consumption was unchanged across the 7 days and showed no effects of strain or sex (*p*>.05 for all).

**Figure 2.**
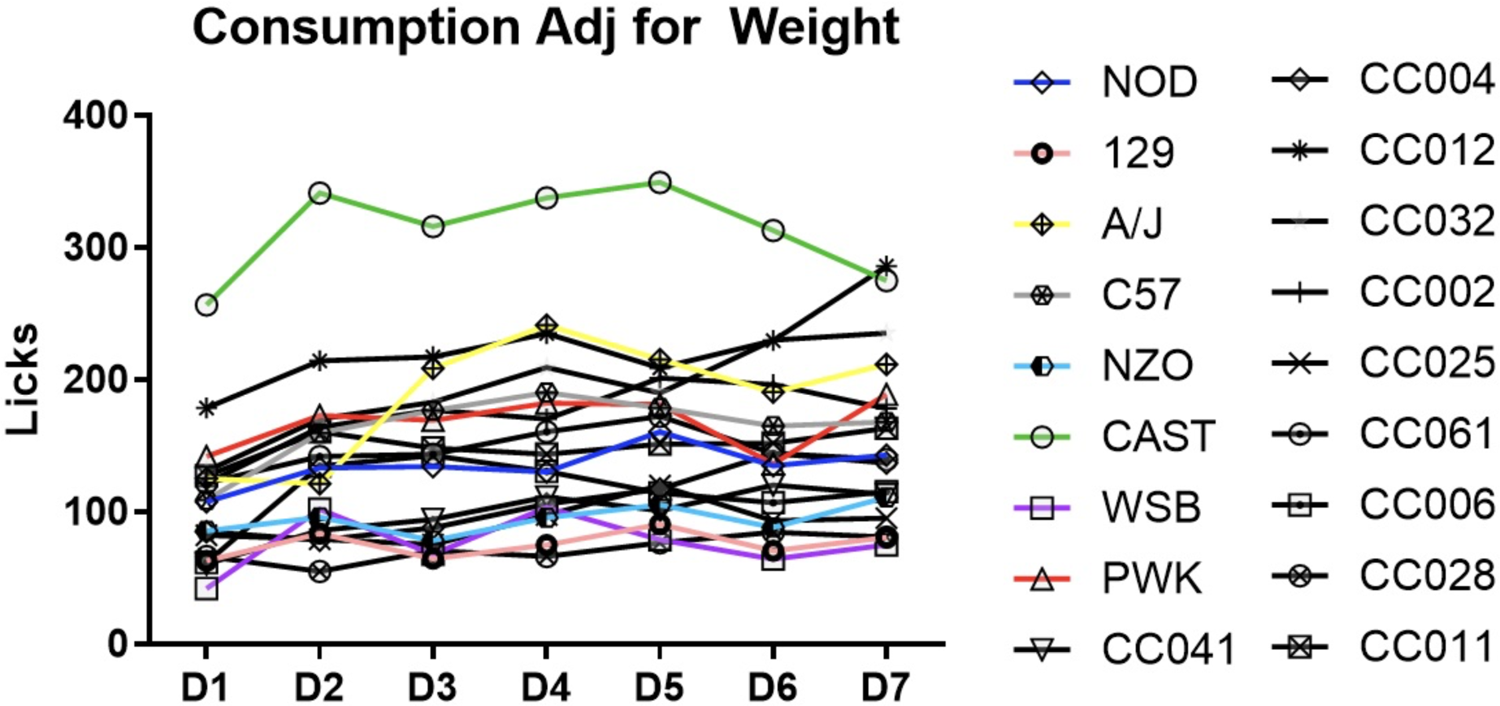
Average total number of Boost licks adjusted for body weight (Licks/g body weight) for each strain daily across seven consecutive days of testing.

A between subjects ANOVA conducted on licks across the final three day average showed a main effect of strain (F[17,278]=14.391, *p*<.001, ηp^2^=.499) and a main effect of sex (F[1, 278]=3.829, *p*=.051, ηp^2^=.015), though no interaction of strain by sex was found (Figure 3). Whether the raw unadjusted or body-weight adjusted licks are examined, CAST/EiJ and A/J mice are the highest licking founder strains, while 129S1/SvImJ and WSB/EiJ are the lowest.

**Figure 3.**
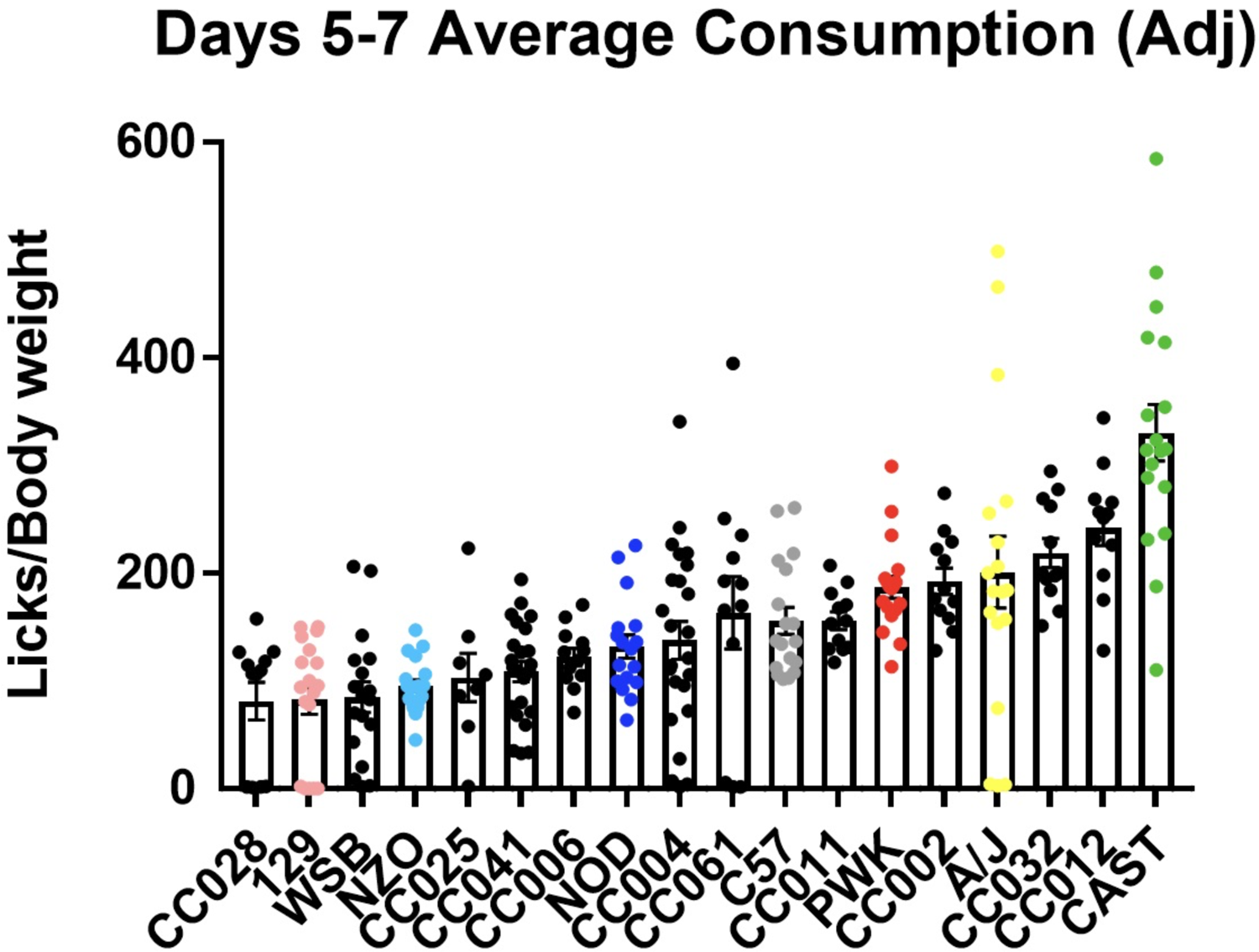
Average of the Boost licks on the last three days of testing (D5-D7) across strains, adjusted for body weight (licks/g body weight).

Finally, a preference score was calculated for each subject by dividing the number of licks on the Boost spout by the number of total licks (Boost + water spouts). This preference score was again averaged across the final three days of testing and analyzed using a one-way ANOVA. No strain, sex, or strain by sex effects emerged (*p*>.05 for all), though this is likely due to a ceiling effect, as all strains exhibited a very high preference for Boost over water.

### Reversal learning

Reversal learning data was collected from partially overlapping strain panels at two sites (Binghamton University and JAX), and a repeated measures ANOVA was conducted on each dependent variable with site as a main effect. We first examined whether site of assessment affected performance. A test stage (acquisition vs. reversal) by site effect was found for the percentage of correct trials (F[1, 319]=8.128, *p*<.01, ηp^2^=.025), and a main effect of site was observed for premature responding in the correct hole in acquisition (F[1, 314]=6.317, *p*<.05, ηp^2^=.022) and premature responding in the incorrect hole in reversal (F[1, 314]=4.832, *p*<.05, ηp^2^=.017), driven by JAX subjects exhibiting higher premature responding overall. As a consequence, site was included as a covariate in analyses of the premature responding variable. No site effect was observed for any other variables (*p*>.05 for all), and site was subsequently dropped from the model for those variables.

A between subjects ANOVA was conducted on the TTC difference score (Figure 3). A main effect of strain was observed (F[44,320]=1.430, *p*<.05, ηp^2^=.115), with no effect of sex or interaction of strain by sex. Further analysis on the founders using a t-test revealed that the 129S1/SvImJ, NZO/HlLtJ, PWK/PhJ, and WSB/EiJ had difference scores significantly greater than 0, indicating that those four strains took significantly more trials to reach criteria in reversal than in acquisition. None of the founder strains showed difference scores significantly less than 0.

Premature responding was analyzed through two separate between-subjects ANOVAs conducted on premature responding in the correct (reinforced) hole at acquisition and incorrect (previously reinforced) hole at reversal. Premature responses in acquisition demonstrated an interaction of strain by sex (F[43, 319]=1.733, *p*<.01, ηp^2^=.194) and a main effect of strain (F[44, 319]=1.555, *p*<.001, ηp^2^=.382; Figure 4). A/J, CC015/UncJ, CC036/UncJ, and CC059/TauUncJ strains were determined to be extreme high premature responders by a Tukey post-hoc comparison, while no extreme lows were detected. Premature responses in reversal showed a main effect of strain (F[44, 320]=3114, *p*<.001, ηp^2^=.300); no sex effects were observed (Figure 5). A/J and CC036/UncJ strains again emerged as an extreme high responder, as did the NZO/HlLtJ and CC061/GeniUncJ. Two wild-derived strains, PWK/PhJ and CAST/EiJ, were found to be extreme lower responders.

**Figure 4.**
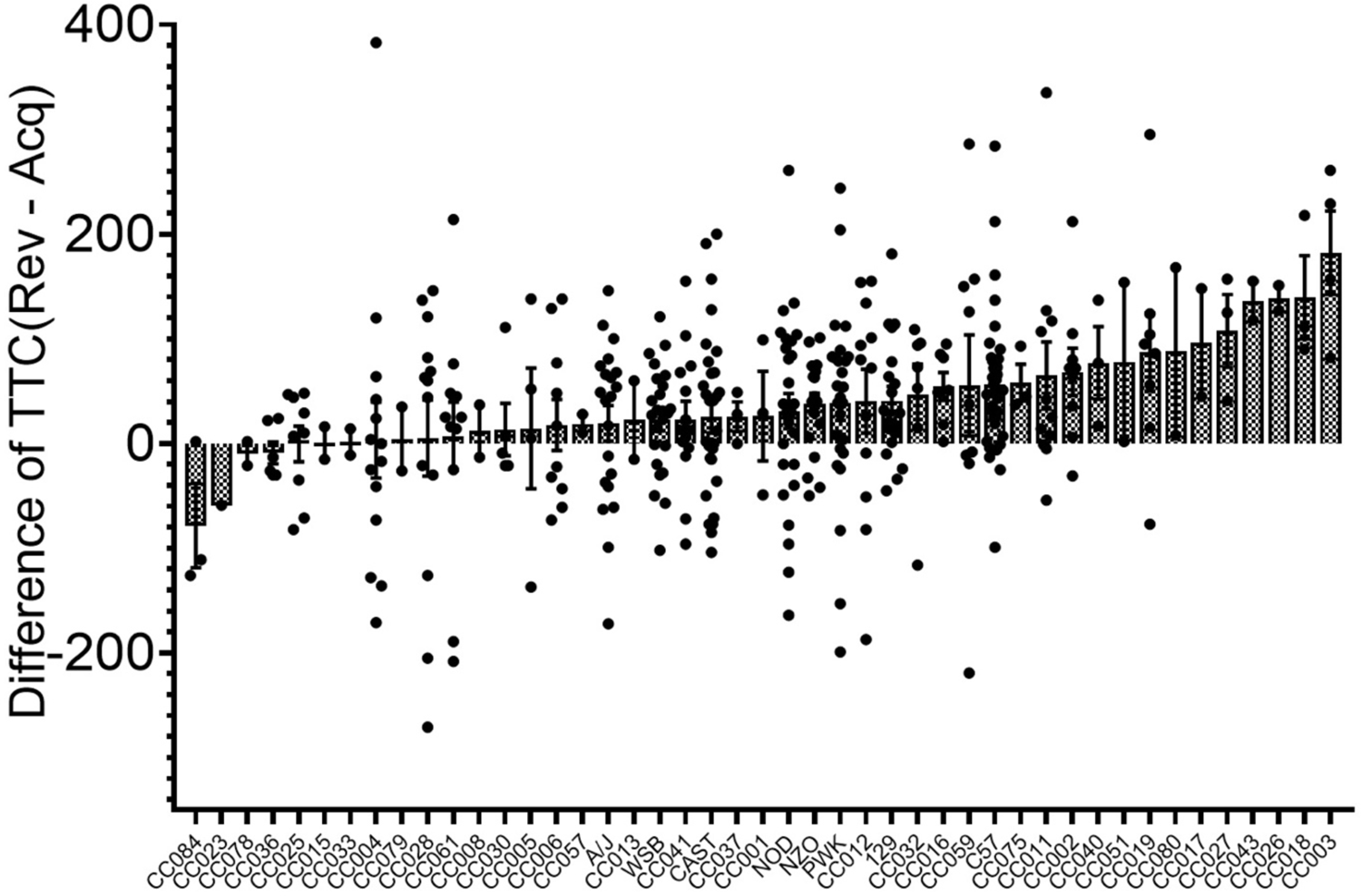
Average total trials in acquisition and reversal stages across strain displayed as a difference score (Reversal-Acquisition). A positive value shows an increase from acquisition to reversal, which demonstrates cognitive inflexibility and impulsive action.

**Figure 5.**
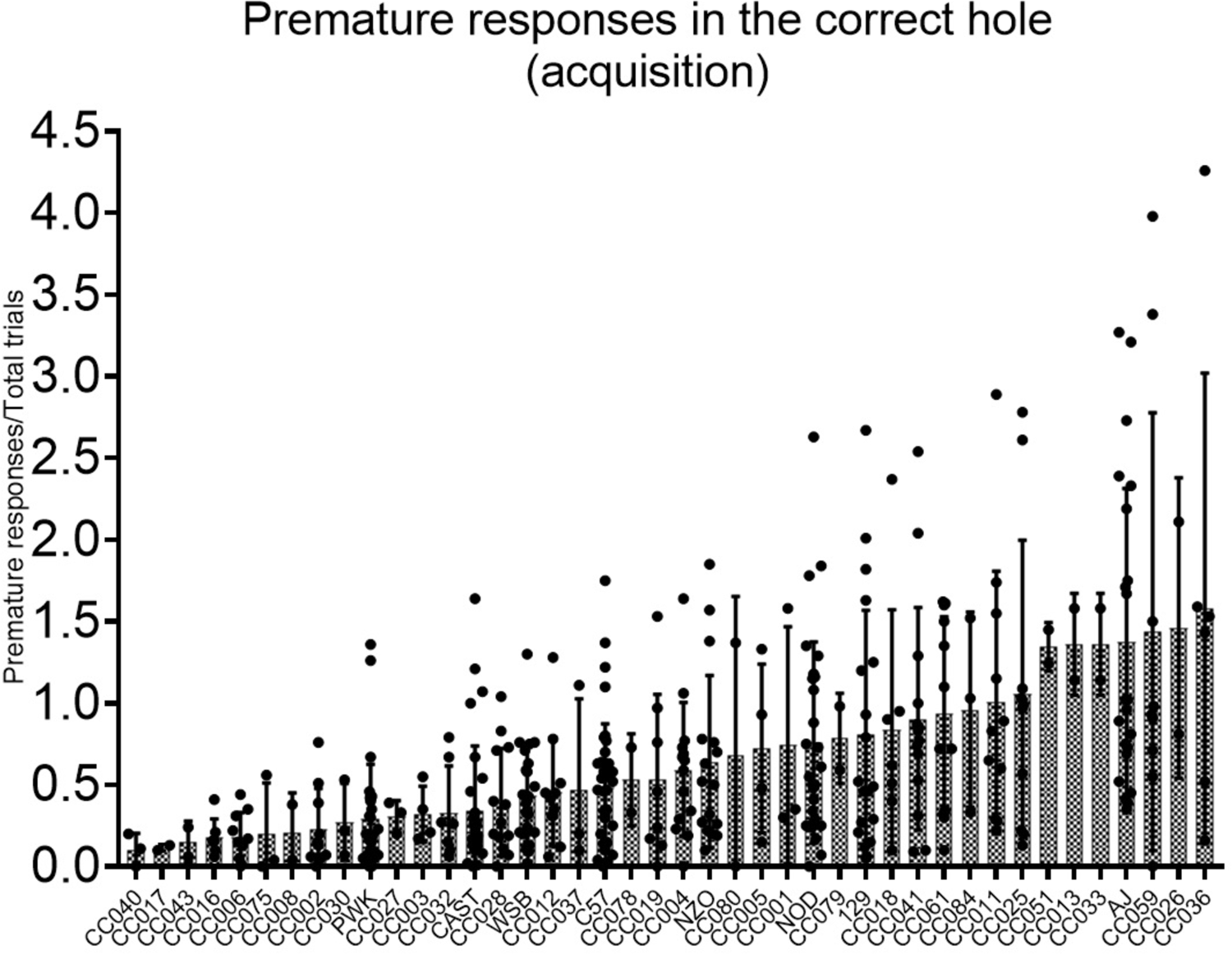
Average premature responses in the correct (rewarded) hole per trial (total premature responses/total trials) in acquisition across strain. A high value represents an inability to withhold responding and high waiting impulsivity.

Ancillary variables were also examined (Table 6) for heritability. Strain impacted the tendency to omit response on trials, as well as the time required to initiate trials and to retrieve earned rewards. None of these strain effects were moderated by sex.

**Table 6.**
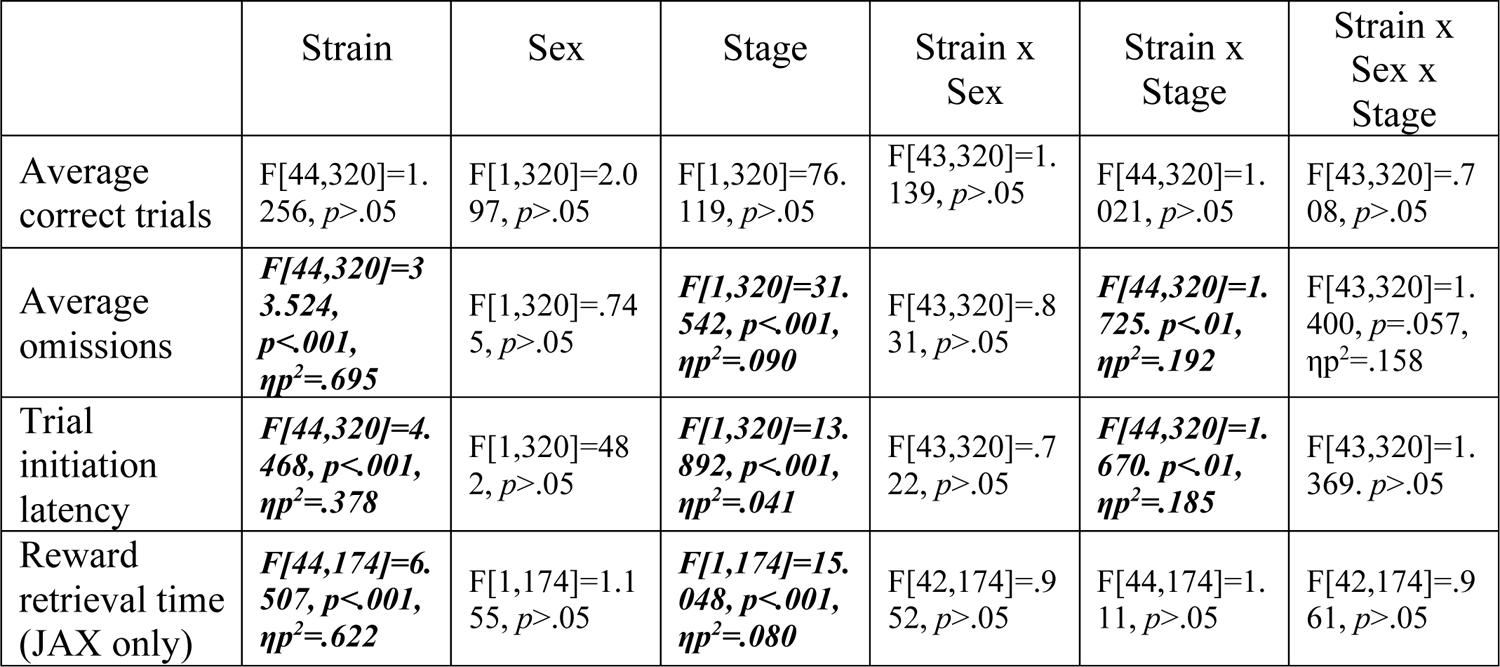
ANOVA statistics for ancillary variables examined in reversal learning. Significant findings are bolded and italicized.

### Delay discounting

A repeated measures ANOVA was conducted on difference score variables as a function of four delay times, revealing a three-way interaction between strain, sex, and delay (F[51, 117]=1.463, *p*<.05, ηp^2^=.175), a trending interaction of strain and delay (F[51, 117]=1.376, *p*=.056, ηp^2^=.167) and expected main effects of delay (F[3, 117]=28.084, *p*<.001, ηp^2^=.194) and strain (F[17, 117]=2.296, *p*<.01, ηp^2^=.250). Further analysis on each founder strain showed significant discounting in the NOD/ShiLtJ, A/J, C57BL/6J, CAST/EiJ, and PWK/PhJ strains, while the 129S1/SvImJ, NZO/HlLtJ, and WSB/EiJ did not exhibit significant discounting under these test conditions (Figure 7).

**Figure 6.**
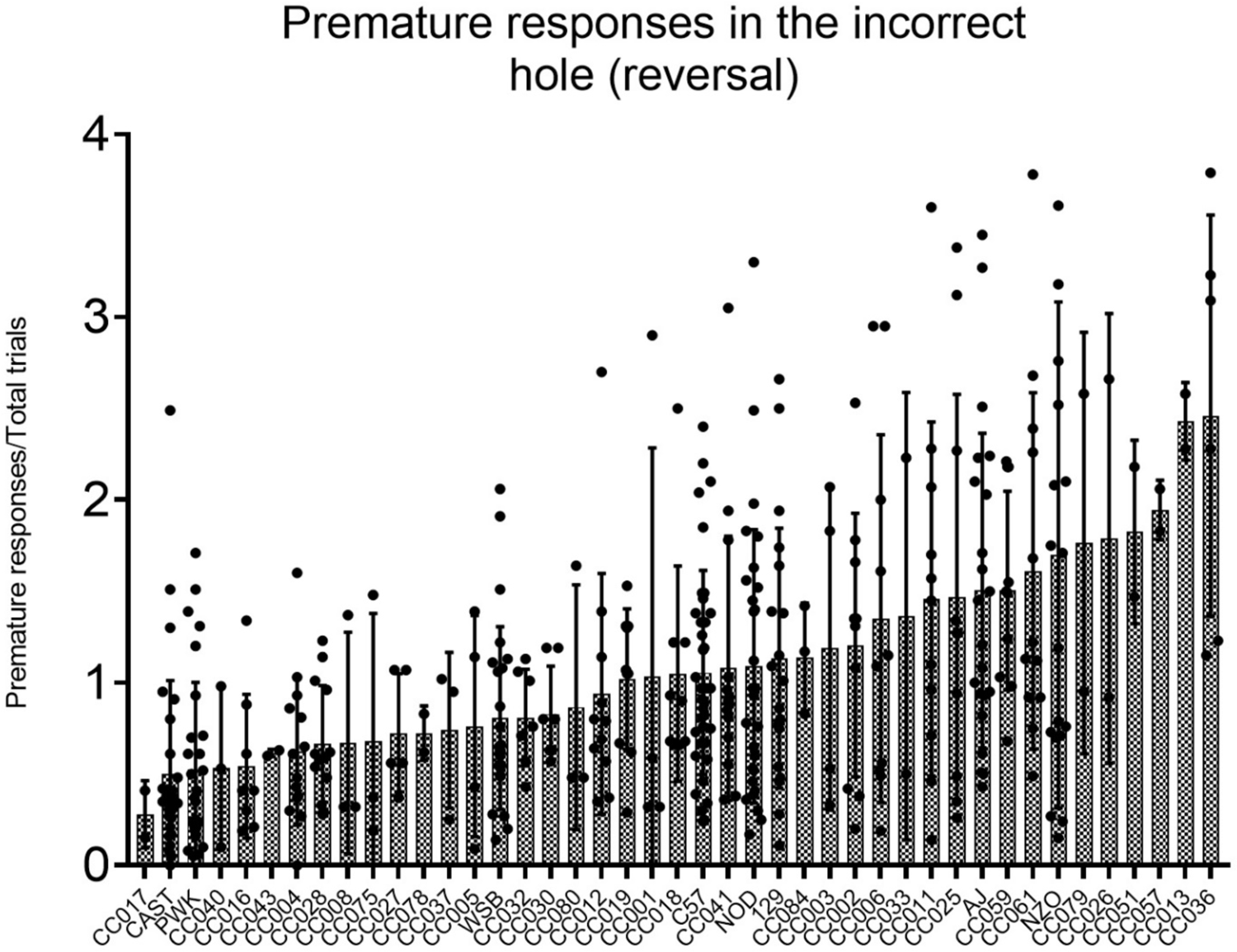
Average premature responses in the incorrect (previously rewarded but currently unrewarded) hole per trial (total premature responses/total trials) in reversal across strain. A high value represents an inability to withhold responding in a previously rewarded hole and high waiting impulsivity.

**Figure 7.**
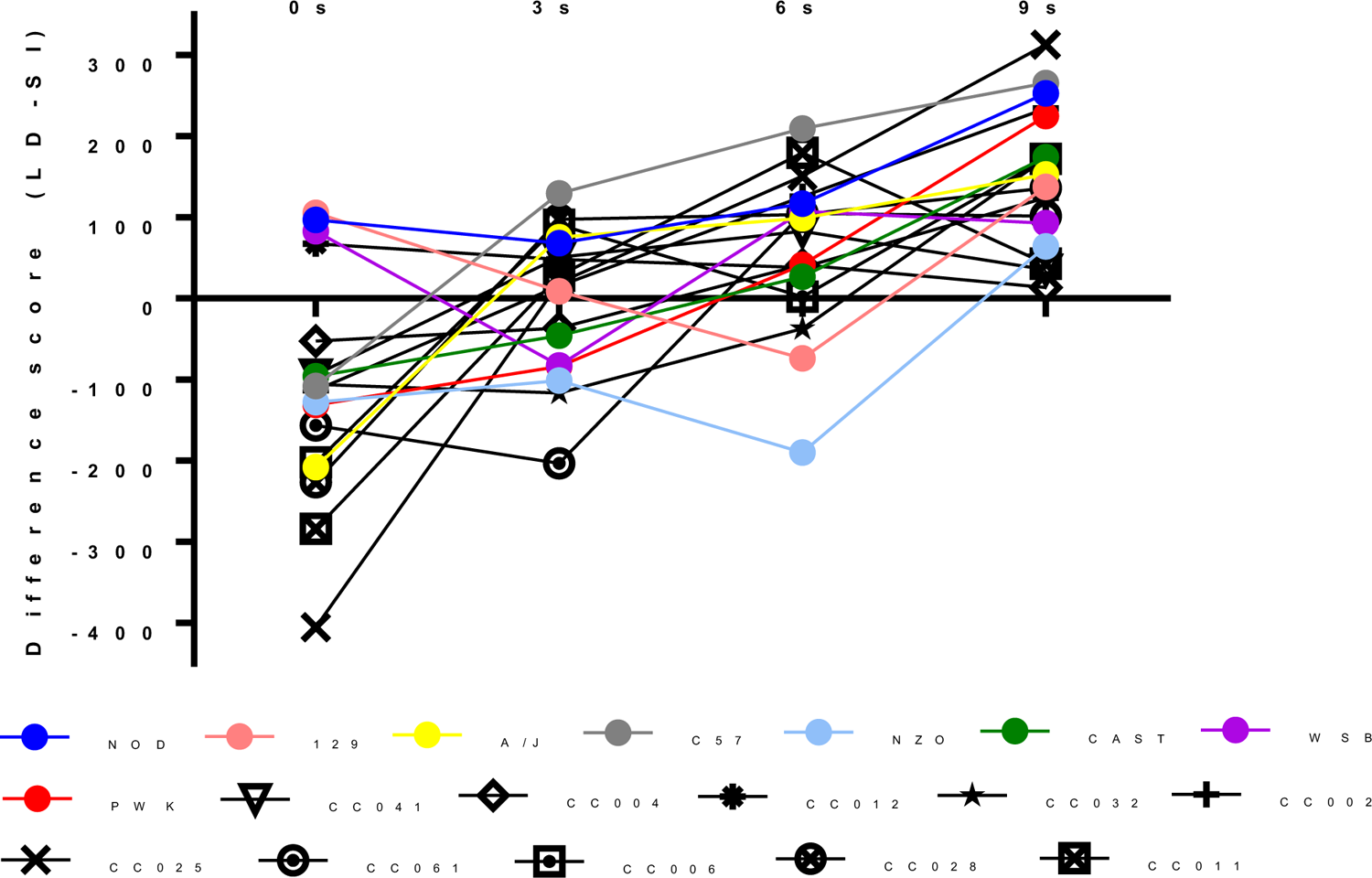
The large delayed (LD) – small immediate (SI) reward amounts at the different delays in delay discounting. A more positive value indicates the LD value was discounted more, demonstrating the SI picked more often and the LD reward value increasing in response. Therefore strain with a steeper slope across the delays more readily discount and show increased impulsive choice, whereas animals with a flatter slope do not demonstrate high discounting behavior.

### Phenotypic correlations

As several studies and groups have used the founder strains in tests of ambulatory response in an open field, we first asked whether our locomotor activity data reproduced existing data. Total ambulatory distance traveled was positively correlated with measures of activity in an open field gathered in other published, publicly available datasets, including Geuther et al., 2019 (MPD(Mouse Phenome Database #):650; *r*[4]=.956, *p*<.001), Kollmus et al., 2020 (MPD:550; *r*[6]=.762, *p*<.01), Kliethermes & Crabbe, 2006 (MPD:599; *r*[6]=.986, *p*<.001), and Wiltshire et al., 2015 (MPD:214; *r*[6]=.951, *p*<.01). Rearing behavior in our dataset was also correlated with that reported by Kliethermes & Crabbe (*r*[6]=.785, *p*<.05) and Wiltshire et al. (*r_s_*[6]=.886, *p*<.05).

Correlations were run on key variables selected from each behavioral test. *A priori* correlations are bolded to show a predicted relationship (Table 7). Locomotor response (distance traveled) to novelty and impulsivity have a conflicting relationship in the literature, with an early study reporting hyperactivity was correlated with impulsive choice (Isles et al., 2004), while later studies found no relationship between activity and impulsive premature responding (Belin et al., 2008; Loos et al., 2009) or delay discounting (Perry et al., 2005). Additionally, the facets of impulsive behavior are dictated by different neural circuits and are thought to reflect uncorrelated phenotypes (Barrus et al., 2015; Dougherty et al., 2009; Nautiyal et al., 2017; Reynolds et al., 2008). However, multiple facets of impulsivity have been identified as having a relationship with over-eating in humans, including non-planning impulsivity, attentional impulsivity, and motor impulsivity (Garza et al., 2016; Georgii et al., 2017; Maxwell, Gardiner, & Loxton, 2020). This relationship has also been observed within rodents as well, with high impulsive rats engaging in more “binge-like” and excessive consumption of highly palatable foods (Anastasio et al., 2019; Velázquez-Sánchez et al., 2014), potentially due to convergent neurocircuitry of the ventromedial prefrontal cortex (Anastasio et al., 2019). A positive correlation is therefore hypothesized between the dimensions of impulsivity and Boost intake.

**Table 7.**
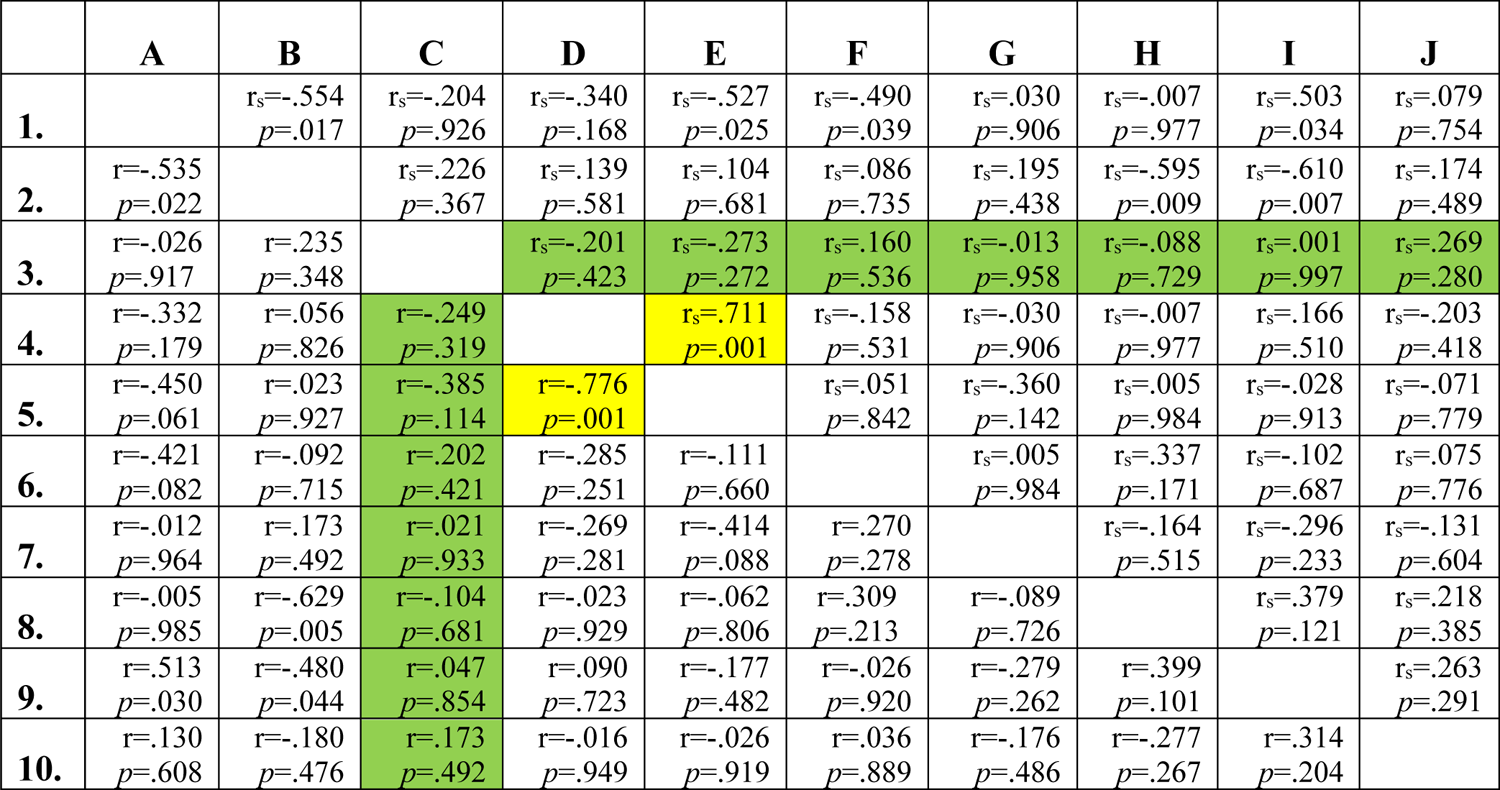
Correlation table for strains collapsed by sex. *A priori* correlations are highlighted in green and significant *a priori* correlations are highlighted in yellow. Spearman correlations are reported on the upper right and Pearson correlations are reported on the lower left. The variables are as follows:

1. Ambulatory distance on D1 of locomotor
2. Ambulatory distance difference score (D2-D1)
3. Boost licks averaged over the final 3 days adjusted for body weight
4. Average premature responses in the correct hole in acquisition
5. Average premature responses in the incorrect hole in reversal
6. Total trial to criteria difference score (Reversal-Acquisition)
7. LD-SI amount at 0s
8. LD-SI amount at 3s
9. LD-SI amount at 6s
10. LD-SI amount at 9s

No strain-level relationship was found between Boost intake and TTC difference score (*r*[16]=.202, *p*=.421), premature responses in the correct hole at acquisition (*r*[16]=-249, *p*=.319) premature responses in the correct hole at reversal (*r*[16]=-.385, *p*=.114), or any of the delay time difference scores (*r*[16]=.021-.173, *p*>.05 for all).

Rate of premature responding during the acquisition and reversal stages was highly correlated (*r_s_*[16]=.711, *p*<.001). While strain-level heritability was identified for each impulsivity measure, no genetic correlation was found between TTC and premature responding (reversal learning) or LD-SI reward difference at any of the delays (delay discounting), suggesting that they are influenced by from distinct genetic architectures (*r*[16]=-.203-.166, *p*>.05 for all).

A Pearson’s correlation was additionally run with key dependent variables and variables of interest utilizing the CC mice and their founders in separate addictions-relevant studies. No relationship was found between ambulation, palatable food reward intake, or impulsivity with measures of ethanol intake (*r*[16]=-0.359—0.209, *p*>.05 for all) or preference (*r*[16]=0-302— 0.130, *p*>.05 for all) in a two bottle choice modified drinking in the dark paradigm (Bagley & Jentsch, 2020), utilizing the same eighteen strains.

Finally, we correlated locomotor and impulsivity measures to acoustic startle response (ASR) and pre-pulse inhibition (PPI) data reported in a separate study (Kollmus et al., 2020) in the founder strains. Impulsive phenotypes and ASR/PPI are dependent on ventrostriatal dopamine transmission, and a correlation between impulsivity and impaired PPI has been identified within human literature (Gee et al., 2015; Swann et al., 2013). No relationship emerged between impulsivity and ASR or PPI, though a strong negative correlation was found between ASR (120DB) and degree of habituation for rearing (*r*[6]=-817, *p*<.05) and stereotypy (*r*[6]=-.957, *p*<.001). This correlation demonstrates that a heightened acoustic startle response appears to be correlated in strains that do not habituate rearing and stereotypy behaviors upon repeated exposure to a novel environment.

## Discussion

The goal of this study was to examine a panel of CC strains and CC founders for heritable addiction-related traits and further understand the relationship between different components of impulsivity. We have identified multiple heritable phenotypes in the founder strains and the ten CC strains, specifically locomotor activity in response to novelty, degree of habituation within the open field apparatus, reward sensitivity as measured by total Boost consumption, aspects of behavioral flexibility and impulsivity measured in reversal learning, and impulsive choice in delay discounting. Overall our results demonstrate a genetic component of these phenotypes, with varying degrees of heritability.

### Locomotor response

Multiple facets of behavior measured in the open field were found to be heritable across the founders and CC strains. Distance traveled in a novel environment, as well as the degree of habituation of this response during a second test, were both observed to be heritable (heritability of 0.404 and 0.268, respectively), as were measures of the duration of ambulation, resting, stereotypy, and rearing (heritability between 0.452 to 0.695 for all measures). These results are consistent with prior reports, as heritability of related phenotypes is well-established within the literature on inbred mouse models (Jeste et al., 1984; Mhyre et al., 2005; Swallow et al., 1998).

As previously discussed, ambulatory distance traveled in response to novelty has a conflicting relationship with impulsivity in the literature. Isles et al. (2004) reported exploratory activity to covary with impulsive choice in four inbred mouse strains. This relationship has been described as impulsive activity, a trait with relevance to attention deficit/hyperactivity disorder (Itohara, Kobayashi, & Nakashiba, 2015). However, subsequent studies have supplied evidence that these two phenotypes are not co-heritable in various strains of laboratory rats (Belin et al., 2008; Loos et al., 2009; Perry et al., 2005), with Perry et al. specifically examining delay discounting, the measure of impulsivity utilized by Isles et al. Ultimately, the lack of a correlation between any traits of ambulation and impulsive behavior in this study support the previous findings that ambulation and impulsivity are disassociated traits (Loos et al., 2009), though this relationship may also be a distinction between rats and mice, or otherwise dependent on the subjects’ genetic background.

### Palatable food intake

The strains were phenotyped for intake of a highly palatable food, providing essential information about hedonic responses to that food and/or reward sensitivity. Past research has suggested sensitivity to reward is a trait that predicts motivation to seek out reinforcing stimuli (Davis & Fox, 2008), and presentation of food and drug cues result in activation of similar regions, as well as activation of similar gene expression patterns (Kelley, Schiltz, & Landry, 2005). Moderate heritability estimates were found for palatable food intake (estimates of .499 and .541 for overall average intake or last three days, respectively). No genetic relationship was found between palatable food intake and impulsivity in the strains of the current study. Human studies have demonstrated that food cravings are a component of trait impulsivity (Garza et al., 2016; Georgii et al., 2017; Nederkoorn et al., 2009), perhaps due to convergent circuitry in the dorsolateral prefrontal cortex (Georgii et al., 2017). This is further supported by a study that segregated Sprague-Dawley rats based on motor impulsivity (determined by the 1-CSRTT), finding that highly impulsive rats engaged in more vigorous binge-like consumption of a high-fat food and that activation of the ventromedial frontal cortex suppressed both impulsive and binge-like feeding behaviors (Anastasio et al., 2019). However, at least one mouse study has found the opposite relationship. When Warthen et al. (2016) stimulated pyramidal neurons in the medial frontal cortex of mice, operant responding for food rewards was increased, while impulsive responding was suppressed. Importantly, this study also found no effect on consumption of freely available food. It is likely that the interaction between trait impulsivity and consumption behaviors, including reward sensitivity, is dependent on a myriad of factors, including internal states such as mood (Herman et al, 2018; Tice et al., 2001) and interoception (Herman et al., 2018; Herman et al., 2019). Nevertheless, results from the current study support the idea that impulsivity phenotypes and consumption of a palatable food reward do not covary in the CC population.

### Impulsivity

Impulsive action and choice were characterized in all strains and found to be significantly heritable for all outcomes which included: reversal learning (0.115), premature responding (0.382 and 0.300 for acquisition and reversal, respectively), and discounting of delayed rewards (0.250). This supports previous findings in both human and animal literature that impulsivity is a heritable trait. Heritability estimates for premature responding reported here are greater than past estimates derived from measures of response to selection in outbred rats (Jupp et al., 2020). Delay discounting heritability estimates are on par with estimates that have been reported previously for inbred mice populations (Isles et al., 2004; Wilhelm & Mitchell, 2009), while estimates for behavioral flexibility in reversal learning are lower than what has been reported in inbred BXD mice (Laughlin et al., 2011). Heritability estimates are no doubt influenced by the specific species and population being evaluated.

The results of this study support the theory proposed by Evenden (1998) that dimensions of impulsivity are dissociable. We found no correlations between impulsivity-relevant phenotypes measured during reversal learning or delay discounting. While each of the types of impulsivity tested do share some mechanistic basis within the frontal cortex and striatal circuitry, each trait demonstrates unique patterns of regional activation. Reversal learning requires function of the orbitofrontal cortex and dorsomedial striatum for optimal inhibitory control (Boulougouris, Dalley, & Robbins, 2007; Chudasama & Robbins, 2003; Clarke, Robbins, & Roberts, 2008; Jentsch et al, 2014), also demonstrated by a functional neuroimaging study in human adults (Ghahremani et al., 2010). Conversely, tests of impulsive choice rely less robustly on the orbitofrontal cortex and are more strongly associated with activation of the lateral frontal cortex (Cho et al., 2010; Hinvest et al., 2011) and the hippocampus (Abela & Chudasama, 2012; Cheung & Cardinal, 2005; Jentsch et al., 2014; McHugh et al., 2008). Waiting impulsivity, by contrast, is most dependent upon activation of the infralimbic cortex, nucleus accumbens, and the subthalamic nucleus (Baunez et al., 1995; Chuadasama et al., 2003; Jentsch et al., 2014; Morris et al., 2016; Voon, 2014). Altogether, these three types of impulsivity seem to be distinct phenotypes that arise from cortico-striato-thalamo dysfunction, which are in part influenced by genetics.

### Inter-lab replicability

Separate reversal learning testing at two sites (Binghamton and JAX) showed that majority of variables analyzed did not have a site effect and therefore demonstrated strong replicability. The site-dependent effects were observed for correct trials and premature responding, the latter of which was driven by JAX site mice demonstrating higher premature responding overall. While each site followed identical testing procedures, Binghamton mice were tested on reward sensitivity and open field prior to reversal learning, while JAX mice were first tested on open field, light/dark, hole board, and novelty place preference. The site effect only on select variables also indicates that certain phenotypes may be uniquely affected by adolescent shipping and separate site variability. Suppressed premature responding at the Binghamton site is a unique finding that cannot definitely be attributed to any factor between the sites. Further research is needed to understand the potential relationship between motor impulsivity and factors such as adolescent shipping stress despite a lengthy habituation period.

### Limitations

Single housing was utilized during testing due to high levels of aggression observed in some strains. Group housing was utilized upon initial acclimation to minimize shipping and acclimation stress and because aggression was observed more often once food restriction began. Evidence for an effect of single housing on impulsivity is mixed. One study isolated rats at PND 28 and found no difference on impulsivity in the 5-CSRTT at adulthood, though did note that these isolated animals were slower to collect food rewards (Dalley et al., 2002). Interestingly, a separate study examined the effects of isolation rearing on delay discounting and noted that isolation reared rats showed reduced impulsive action and impulsive choice (Liu, Wilkinson, & Robbins, 2017). With this in mind, it also must be taken into consideration that certain strains may be more sensitive to the effects of isolation housing, and the degree to which operant performance is altered may vary.

A small amount of caffeine is present in the Boost reinforcer and may exert an effect on animals tested at the Binghamton site, particularly if one or more strains are exceptionally sensitive to it. Chocolate Boost contains .62 mg of caffeine per fluid ounce (Caffeine Informer). Each reward delivery is approximately 20 μl, or .00068 fluid ounces, resulting in mice receiving .00418 mg caffeine per reinforcer. The maximum reinforcers a mouse received on average was 80 in delay discounting, with less rewards being received in reversal learning. Thus, a mouse could possibly receive at maximum of .3344 mg of caffeine per day. This was converted into a mg/kg dose for a low weight animal (12g) and a high weight animal (40g). Respectively the daily dose was calculated to be 10 mg/kg and 3 mg/kg (PO). A bolus dose of 15 mg/kg i.p. caffeine is described as being a moderate dose (Hnasko, Sotak, & Palmiter, 2005), and 1.5 mg/kg i.p. doses were found to be enough to produce conditioned place preference in mice (Patkina & Zvartau, 1998). Past studies have shown that .5-16 mg/kg i.p. dose of caffeine increases locomotor activity in mice (Kayir & Uzbay, 2004), and i.p. doses of 5mg/kg-15mg/kg increase wakefulness (Huang et al., 2005). Route of administration additionally plays a factor: oral consumption of caffeine decreased the amount of cocaine later self-administered, while 3 mg/kg i.p. injections increased it despite similar metabolite levels (Kuzmin et al., 2000). This information indicates that mice in the present study were receiving variable levels of caffeine that depended on number of reinforcers received, though this was at maximum a low-moderate dose, and consumption was not impacted by body weight.

## Conclusion

Altogether, the present study marks one of the first systematic attempts to phenotype the CC founder strains for ambulatory activity, reward sensitivity, and three types of impulsivity. Heritability for these traits has been identified, and the lack of genetic correlation between each of these types of impulsivity implicate unique neurogenetic factors between traits. These results suggest that genetically diverse populations like the CC used in this study and Diversity Outbred (DO) populations are suited for forward genetic approaches to reveal unique genetic factors that influence dimensions of impulsivity and how these unique factors influence risk for SUDs.

